# A regularized functional regression model enabling transcriptome-wide dosage-dependent association study of cancer drug response

**DOI:** 10.1101/2020.06.18.158907

**Authors:** Evanthia Koukouli, Dennis Wang, Frank Dondelinger, Juhyun Park

## Abstract

Cancer treatments can be highly toxic and frequently only a subset of the patient population will benefit from a given treatment. Tumour genetic makeup plays an important role in cancer drug sensitivity. We suspect that gene expression markers could be used as a decision aid for treatment selection or dosage tuning. Using *in vitro* cancer cell line dose-response and gene expression data from the Genomics of Drug Sensitivity in Cancer (GDSC) project, we build a dose-varying regression model. Unlike existing approaches, this allows us to estimate dosage-dependent associations with gene expression. We include the transcriptomic profiles as dose-invariant covariates into the regression model and assume that their effect varies smoothly over the dosage levels. A two-stage variable selection algorithm (variable screening followed by penalised regression) is used to identify genetic factors that are associated with drug response over the varying dosages. We evaluate the effectiveness of our method using simulation studies focusing on the choice of tuning parameters and cross-validation for predictive accuracy assessment. We further apply the model to data from five *BRAF* targeted compounds applied to different cancer cell lines under different dosage levels. We highlight the dosage-dependent dynamics of the associations between the selected genes and drug response, and we perform pathway enrichment analysis to show that the selected genes play an important role in pathways related to tumourgenesis and DNA damage response.

**Author Summary:** Tumour cell lines allow scientists to test anticancer drugs in a laboratory environment. Cells are exposed to the drug in increasing concentrations, and the drug response, or amount of surviving cells, is measured. Generally, drug response is summarized via a single number such as the concentration at which 50% of the cells have died (IC50). To avoid relying on such summary measures, we adopted a functional regression approach that takes the dose-response curves as inputs, and uses them to find biomarkers of drug response. One major advantage of our approach is that it describes how the effect of a biomarker on the drug response changes with the drug dosage. This is useful for determining optimal treatment dosages and predicting drug response curves for unseen drug-cell line combinations. Our method scales to large numbers of biomarkers by using regularisation and, in contrast with existing literature, selects the most informative genes by accounting for drug response at untested dosages. We demonstrate its value using data from the Genomics of Drug Sensitivity in Cancer project to identify genes whose expression is associated with drug response. We show that the selected genes recapitulate prior biological knowledge, and belong to known cancer pathways.

## Introduction

Cancer is a heterogeneous disease, with individual tumours showing sometimes very different mutational and molecular profiles. The genetic makeup of a tumour influences how it reacts to a given anti-cancer drug. However, due to lack of predictive markers of tumour response, often patients with very different tumour genetic makeup will receive the same therapy, resulting in high rates of treatment failure [1]. Large clinical trials in rapidly lethal diseases are expensive, complex and often lead to failure due to lack of efficacy at a given dosage [2]. One major issue for some cancer treatments, e.g. chemotherapies, are cytotoxic effects that result in collateral damage of the healthy host tissue [3]. Patient remission depends not only on the selection of the right drug but also on the determination of the optimal dosage, especially when drugs with small therapeutic range, high toxicity levels or both are administered. Genetic factors can help fine-tune the dosage for individual patients, so that the minimal effective dosage can be delivered [4].

Treatment response in patients with specific cancers had been intensely examined in relation to the molecular characteristics of the tumours [5]. However, cellular heterogeneity within the tumour and the lack of standard metrics for quantifying drug response in patients can make it difficult to computationally model response as a function of molecular features. Cancer cell line drug screens can provide valuable information about the effect of genetic features on drug dose-response in a controlled setting. During the last decade, there have been several systematic studies that examined the relationship between genetic variants and drug response in cell lines [6–10]. There have also been studies that measured transcriptional profiles [11,12] and drug response in cancer cells after administering anticancer drugs at various dosages [13, 14]. By comparing multiple genomic features of cell lines to drug response, the investigators were able to identify gene signatures for drug responsiveness in specific cancer types. However, these signatures were selected based on a single summary statistic of response, usually IC50, that may not always be the most useful metric for differentiating drugs [15], and only provides information on one dose concentration. While these existing signatures of drug response provide a way towards selecting the right drug for a patient, none of them characterise gene-dose relationships that may ultimately identify the optimal dose for a drug to use in the clinic.

With regards to the high-dimensional nature of genomic datasets, it is worth noting that highly-complex data sets with non-stationary trends are not easily amenable to analysis by classic parametric or semi-parametric mixed models. Such effects, e.g. the effect of genes on drug response over different drug dosages (dose-varying effect), can be examined using varying coefficient models which allow for the covariate effect to be varying instead of constant [16]. Methods to estimate varying covariate effects include global and local smoothing, e.g. kernel estimators [17, 18], basis approximation [19] or penalised splines [20]. Although non-parametric techniques can reduce modelling biases [21], they often suffer from the “curse of dimensionality” [22]. Inference in these models becomes impossible as the number of predictors increases, and often selecting a smaller number of important variables for inclusion into the model is clinically beneficial. Sparse regression has enabled a more flexible and computationally “inexpensive” way of choosing the best subset of predictors [23]. However, these methods cannot handle ultra-high dimensional problems without losing statistical accuracy and algorithmic stability, since they handle all of the predictors jointly. Consequently, there is a need of prior univariate tests focused on filtering out the unimportant predictors by estimating the association of each predictor to the outcome variable separately [21,24,25]. The advantage of using varying coefficient models along with a variable screening algorithm on genomic data sets was first introduced to explore the effect of genetic mutations on lung function [24]. Recently, Tansey et al. [26] proposed a method for modelling drug-response curves via Gaussian processes and linking them to biomarkers using a neural network prediction model. The authors did not use their model for dosage-dependent inference of biomarker effects, and the highly non-linear neural network model makes interpretation of biomarker effects challenging.

Here, we extended the methodology of Chu et al. [24] to the objective of assessing the transcriptomic effect on anti-cancer drug response, where our coefficient functions were allowed to vary with dosage. We developed a functional regression framework to study the effectiveness of multiple anticancer agents applied in different cancer cell lines under different dosage levels, adjusting for the transcriptomic profiles of the cell lines under treatment. We considered a dose-varying coefficient model, along with a two-stage variable selection method in order to detect and evaluate drug-gene relationships, and then applied this method to data extracted from the Genomics for Drug Sensitivity in Cancer (GDSC) project [7]. To compare and differentiate similar treatments, we examined a case study of five BRAF targeted compounds under different dosages to almost 1000 cancer cell lines. We used baseline gene expression measurements for the cancer cell lines to investigate gene-drug response relationships for almost 18000 genes. Gene rankings were obtained based on the estimated effects of the genes on the drug response. The resulting model describes the whole dose-response curve, rather than a summary statistic of drug response (e.g. IC50), which allowed us to identify trends in the gene-drug association at untested dose concentrations.

## Materials and Methods

### The Genomic of Drug Sensitivity in Cancer data

Drug sensitivity data and molecular measures derived from 951 cancer cell lines used for the screening of 138 anticancer compounds were downloaded from the GDSC database (https://www.cancerrxgene.org/). We specifically focused on cell lines of cancers of epithelial, mesenchymal and haematopoietic origin treated by five BRAF targeted inhibitors (PLX-4720, Dabrafenib, HG6-64-1, SB590885 and AZ628; GDSC1 data). The maximum screening concentration for each different drug was: 10.00 uM for PLX-4720 and Dabrafenib, 5.12 uM for HG6-64-1, 5.00 uM for SB590885 and 4.00 uM for AZ628. Additionally, we used the independently generated GDSC2 dataset to validate our approach on drugs targeting *MEK1, MEK2* genes (Trametinib-1.00 uM; Selumetinib-10.00 uM, and; PD0325901-0.250 uM) and the PI3K/MTOR signalling pathway (Alpelisib-10.00 uM; AMG-319-10.00 uM, and; AZD8186-10.00 uM). The drug sensitivity measurement was obtained via fluorescence-based cell viability assays 72 hours after drug administration [7]. Approximately 66% of drug sensitivity responses were measured over nine dose concentrations (2-fold dilutions) and 34% were measured over five drug concentrations (4-fold dilutions). In total, we considered 3805 cancer cell line-drug combinations (experimental units). The distribution of different tissues of origin treated were similar across the different drugs tested (for additional information see Fig S2). Paired microarray gene expression data (17737 genes) was available together with the drug response dataset (https://www.cancerrxgene.org/gdsc1000/GDSC1000_WebResources/Home.html).

The dose-response dataset also included a blank response for wells on the experimental plate that had not been seeded with cells or treated with a drug. Blank responses have been used to adjust for the magnitude of the observation error while measuring the amount of cells in each well. We used an affine transformation to the reported responses in order to normalise them within the drug concentration interval, 0 (0% of the maximum dosage) to 1 (100% of the maximum dosage). In particular, for the normalising procedure, we have used the formula:

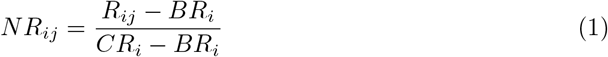

where *R_ij_* is the response of the *i^th^* subject at the *j^th^* dosage level, *CR_i_* is the response under no drug administration (zero dose, *n_i_* = 1), *BR_i_* is the blank response of the *i^th^* subject as described above and *NR_ij_* is the new score taken from the transformation, *i* = 1, …, 3805, *j* = 1, …, *n_i_*.

### A two-stage algorithm for identification of gene-drug associations

Non-parametric techniques have been a great tool for reducing modelling bias and producing data driven inference. Flexible modelling techniques applied on high-dimensional genomic data sets can cause real problems in inference making. Sparse regression techniques, such as the LASSO, can be used as dimensionality reduction techniques, but cannot handle ultra-high dimensional problems without introducing statistical inaccuracies, algorithmic instability and a huge computational burden. Hence, the need for a feature screening algorithm which will marginally filter unimportant variables becomes essential [23]. Below we further explain the two-stage algorithm that has been built in order to detect and explore dose-dependent gene-response associations.

Let the repeated measures data {(*d_ij_*, *y_ij_*, ***z**_i_*, ***x**_i_*) : *j* = 1, …, *n_i_*, *i* = 1, …, *n*}, where *y_ij_* is the response of the *i*th experimental unit (corresponds to a drug sensitivity assay of a specific drug on a specific cell line) at the jth drug dosage level *d_ij_* and ***z**_i_* along with ***x**_i_* are the corresponding vectors of scalar (dose-invariant) covariates. The covariate vector ***z**_i_* = (1, *z*_*i*1_, …, *z_ip_*)*^T^* is a low-dimensional vector of predictors that should be included in the model, whereas ***x**_i_* = (*x*_*i*1_, *x*_*i*2_, …, *z_iG_*)*^T^* is a high-dimensional vector, i.e. 17737 gene expression measurements, that needs to be screened. We assumed that only a small number of *x*-variables (in our case, genes) are truly associated with the response while most of them are expected to be irrelevant; i.e. we make a sparsity assumption.

To explore potential dose-varying effects between the covariates and the drug response, we consider the following varying coefficient model:

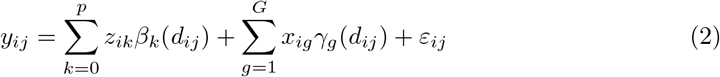

where {*β_k_*(·), *k* = 0, …, *p*} and {*γ_g_*(·), *g* = 1, …, *G*} are smooth functions of dosage level 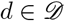, where 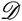 is a closed and bounded interval of 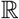. The errors *ε_ij_* were assumed to be independent across subjects and potentially dependent within the same subject with conditional mean equal to zero and variance Var(*ε*) = *σ^2^*(*d*) = *V*(*d*).

Methods for estimating the coefficient functions in Eq (2) include local and global smoothing methods, such as kernel smoothing, local polynomial smoothing, basis approximation smoothing etc. For computational convenience, in this application we used basis approximation smoothing via B-splines.

Let the sets of basis functions {*B_lk_*(·) : *l* = 1, …, *L_k_*} and 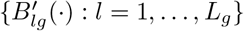 and constants {*ζ_lk_* : *l* = 1, …, *L_k_*} and {*η_lg_* : *l* = 1, …, *L_g_*} where *k* = 0, …, *p* and *g* = 1, …, *G* such that, 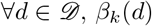 and *γ_g_*(*d*) can be approximated by the expansion

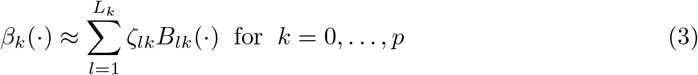

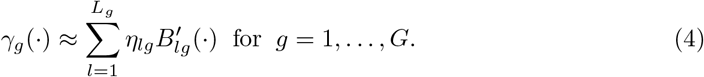

Substituting *β_k_*(·) and *γ_g_*(·) of Eq (2) with Eq (3) and Eq (4), we approximated Eq (2) by

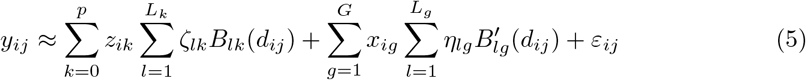

If *B_k_*(·) and 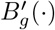 are groups of B-spline basis functions of degree *q_k_* and *q_g_* respectively, and *δ*_0_ < *δ*_1_ < … < *δ_K_k__* < *δ_K_k+1__* and *δ*_0_ < *δ*_1_ < … < *δ_K_g__* < *δ*_*K*_*g*_ +1_ are the corresponding knots, then *L_k_* = *K_k_* + *q_k_* and *L_g_* = *K_g_* + *q_g_*.

Using the approximation Eq (5), the coefficients ***ζ*** = (*ζ*_0_, *ζ*_1_, …, *ζ_p_*)*^T^* and ***η*** = (*η*_1_, *η*_2_, …, *η_G_*)*^T^* can be estimated by minimizing the squared error

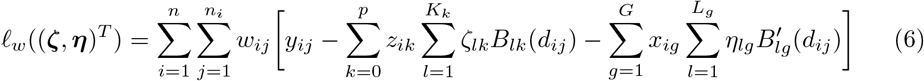

where *w_ij_* are known non-negative weights.

In cases where *p* + *G* >> *n* though, minimisation of Eq (6) is infeasible. Our aim was to identify factors of the covariate vector **x** = (**x**_1_, **x**_2_, …, **x***_G_*)*^T^* (genes) that are truly associated with the response (cancer cell line sensitivity to the drug). In addition, we wanted to explore potential dose-varying effects on the drug response.

We make the following sparsity assumption: any valid solution 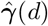 will have 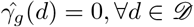 for the majority of components *g*. To detect non-zero coefficient functions we applied a two-stage approach which incorporated a variable screening step and a further variable selection step.

### Screening

The sparsity assumption applies only to components of **x**, the high-dimensional covariate vector in Eq (2).

Let the set of indices

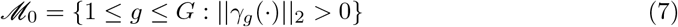

where ‖ · ‖_2_ is the *L*_2_-norm. In order to rank the different components of ***x***, we fitted the marginal non-parametric regression model for the *g*th *x*-predictor:

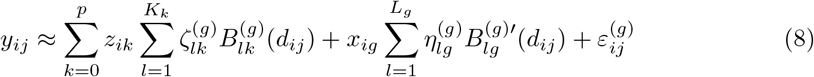

where: 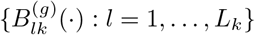 and 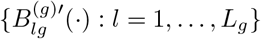 are sets of coefficient functions; 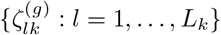 and 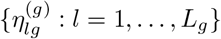 are constants to be estimated, *k* = 0, …, *p*; and, *ε*^(*g*)^ is the error term similar to Eq (5). We then computed the following weighted mean squared error for each *g* ∈ { 1, …, *G*},

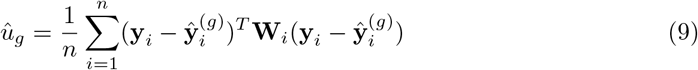

to quantify the importance of the *g*th x-variable. Here,

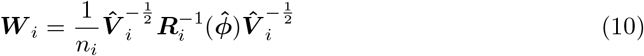

where 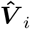 is the *n_i_* × *n_i_* diagonal matrix consisting of the dose-varying variance

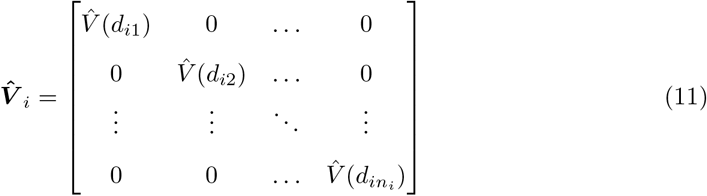

and ***R_i_***(***ϕ***) = (*R_jk_*) the *n_i_* × *n_i_* working correlation matrix for the *i^th^* subject. By ***ϕ***, we denoted the *s* × 1 vector that fully characterises the correlation structure. The estimate of ***ϕ***, 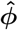, was obtained by taking the moment estimators for the parameters ***ϕ*** in the correlation structure based on the residuals obtained from fitting the following model

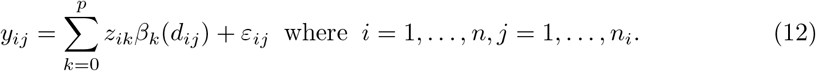

The variance function *V*(*d*) in Eq (11) was estimated using techniques described in [24].

After having obtained 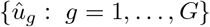, we sorted gene utilities in an increasing order, where smaller 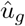 values indicate stronger marginal associations. The *x*-predictors included in the screened submodel are, then, given by

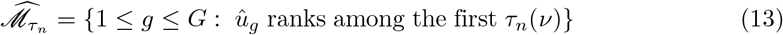

where *τ_n_*(*ν*) corresponds to the size of the submodel which is chosen to be smaller than the sample size *n*.

### Variable selection using a group SCAD (gSCAD) penalty

Screening algorithms aim to discard all unimportant variables but tend to be conservative. In order to preserve only the most important *x*-predictors in the final model, we considered a model including the first *τ_n_*(*ν*) outranked genes and we applied a gSCAD penalty by minimising the following criterion:

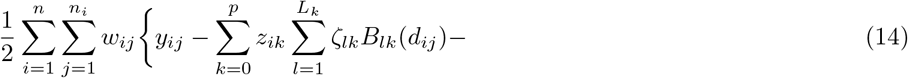

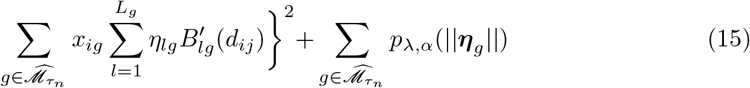

where

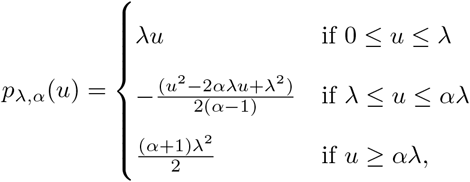

*α* is a scale parameter, λ controls for the penalty size and ‖ · ‖ is the Euclidean *L*_2_-norm. At this point, note that grouping is applied for the coefficients ***η**_g_* that correspond to the same coefficient function. In addition, in order to reduce the bias introduced when applying a LASSO penalty, we alternatively chose the SCAD, which coincides with the LASSO until *u* = λ, then transits to a quadratic function until *u* = *α*λ and then it remains constant ∀*u* > *α*λ, meaning that retains the penalisation and bias rates of the LASSO for small coefficients but at the same time relaxes the rate of penalisation as the absolute value of the coefficients increases. In Fig 1 the reader can find a brief overview of the employed methodology.

**Fig 1.**
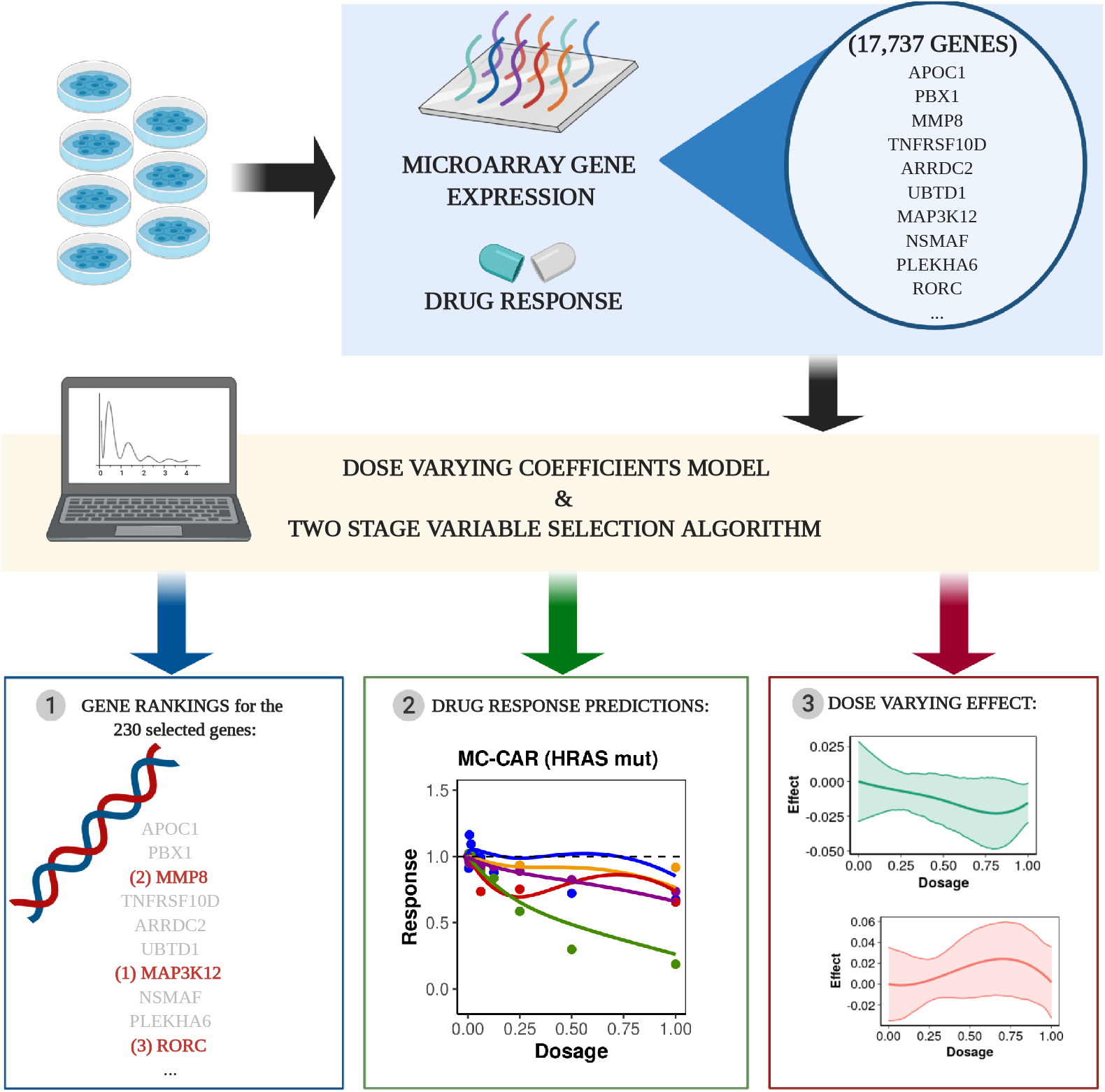
The two-stage algorithm for identifying dose-dependent associations between genes and drugs. Gene expression and drug response data from a drug screening study (e.g. GDSC) are used to fit our dose-varying coefficients model to estimate the dose-varying effect between covariates and drug response. A two-stage variable screening and selection algorithm is applied to rank gene-drug associations. The selected genes can then be used to predict dose-dependent response for drugs of interest.

### Tuning parameters selection

We used knots placed at the median of the observed data values along with cubic B-splines with 1 interior knot, resulting from calculating the number of interior knots suitable using the formula 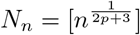 proposed and applied by [19,27,28]. Due to the computational burden this would add, we did not apply cross-validation.

As for the screening threshold *τ_n_*, its magnitude could be determined by the fraction 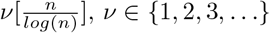. We conducted a pilot simulation study in order to decide the most appropriate size (for further details see Text S1). We also considered an automated algorithm for its selection (Greedy Iterative Non-parametric Independence Screening-Greedy INIS, [29]). Finally, the penalty size for the gSCAD step λ was determined using a 5-fold cross-validation.

### Simulation study

Monte Carlo simulations were conducted to examine the ability of our model to detect the genes that are truly associated with the drug response. This had a key role in tuning model parameters and simultaneously assess model goodness-of-fit using a fraction of the original data set in order to reduce the computational burden of conducting a simulation study under the original dimensions of the data. Responses over different dosage levels were generated based on a subset of genes, the corresponding low-dimensional GDSC data covariates (drug and cancer type) and some specified smooth coefficient functions (see Text S1). Due to the computational burden associated with a simulation of the same scale as the data set, we conducted a simulation study using smaller random fragments of the original GDSC data set. In particular, we repeatedly sampled without replacement 190 experimental units and 886 genes based on which the simulated responses have been generated. The performance of the employed methodology has been assessed based on 1000 simulations using three screening thresholds 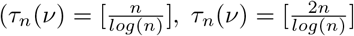 and *τ_n_*(*v*) chosen using the greedy-INIS algorithm [29]) and two estimated covariance structure scenarios (independence and rational quadratic covariance structure). Cubic B-splines and knots placed at the median of the observed data values have been used for estimating the coefficient functions.

To evaluate the performance of the proposed procedure we used the following summary measures: TP–number of genes correctly identified as active; FP–number of the genes incorrectly identified as active; TN–number of the genes correctly identified as inactive; FN-number of the genes incorrectly identified as inactive.

Simulation results suggested that our method accurately detects the drug associated genes from the simulated responses under most of the examined scenarios (Fig A in Text S1). A screening threshold of size 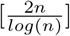 and regression weights adjusted for the covariance structure of the data have been identified as the scenario where our method reached its maximum accuracy. Consequently, for the GDSC application, we chose the screening threshold to be the maximum possible, i.e. 923 genes derived from the formula 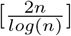, and weights derived by assuming a rational quadratic covariance structure for the repeated measures.

### Data and software availability

The analysis has been conducted using R version 3.6.3. Code for applying the two-stage variable selection algorithm is available online as an R package at https://github.com/koukoulEv/fbioSelect. The data is available online at the Genomics of Drug Sensitivity in Cancer website https://www.cancerrxgene.org/.

## Results and Discussion

### Dose-dependent associations with gene expression in a large-scale drug sensitivity assay

We applied the two stage variable selection algorithm under the dose-varying coefficient model framework described above. Gene rankings and predicted mean drug effects over different dosage levels were obtained. Our algorithm identified 230 candidate genes associated with drug response. The effect of each of those genes was assessed with respect to:

1. the area under the estimated coefficient curve (AUC) and its corresponding standard deviation (estimated using bootstrapping);
2. the effect on cell survival (overall positive, overall negative, mixed);
3. Spearman correlation between the coefficient function value and the dosage level;
4. the mean fold change of the expression of cell lines carrying *BRAF* mutations with respect to wild type; and,
5. the protein-protein interaction network distance between the *BRAF* gene and the selected genes using the Omnipath database [30].

The 230 genes were ranked based on the estimated AUC value (Table S4), and the top 30 genes were highlighted for further analysis (Table 1). The higher the AUC, the larger the effect of the gene on the drug response. The overall effect on cell survival can be either positive, negative or vary over the different dosage levels as determined by the range of the estimated coefficient function. Spearman’s rank correlation was used as an indicator of the coefficient function’s monotonicity by characterising the progress of the genetic effect over different dosage levels. For instance, high expression of the *C3orf58* gene at baseline has a positive effect on cancer cell survival, which becomes stronger as the dosage increases (Spearman’s correlation=0.922). In other words, high expression of this gene can be an indicator of drug resistant cell lines. On the other hand, the *DLC1* gene has a decreasing (Spearman’s correlation=-0.928) and negative effect on cancer cell survival which suggested that as the dosage increases, higher baseline expression of this gene can indicate higher drug sensitivity at higher dosage. Elevated expression of DLC1 has been observed in melanoma and is a well known tumour suppressor that could be a novel marker of BRAF inhibition [31]. Finally, in cases where the overall effect varies (changes between positive and negative), the effect of gene expression on the drug response depends on the drug dosage. In particular, the effect of *DLX6* increases and then decreases at higher dosages (Fig 2). Given the biological and technical variation in drug screens, we should treat the mean effect estimates with caution and consider the confidence intervals of the coefficient functions in order to derive conclusions about the exact effect of the selected genes on the dose response (Fig 2).

**Table 1.**
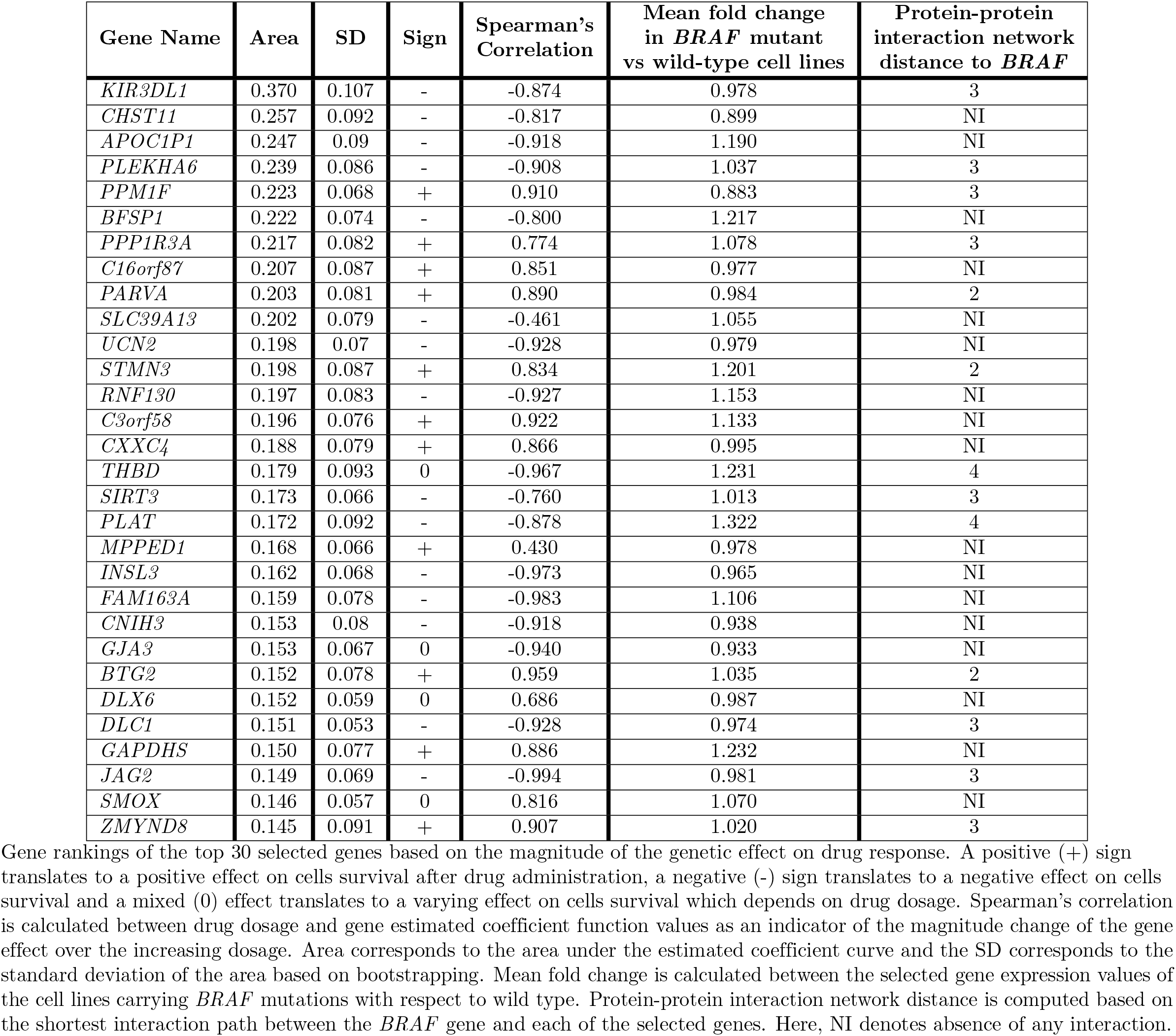
Top 30 gene rankings based on the estimated area under the coefficient function curve.

**Fig 2.**
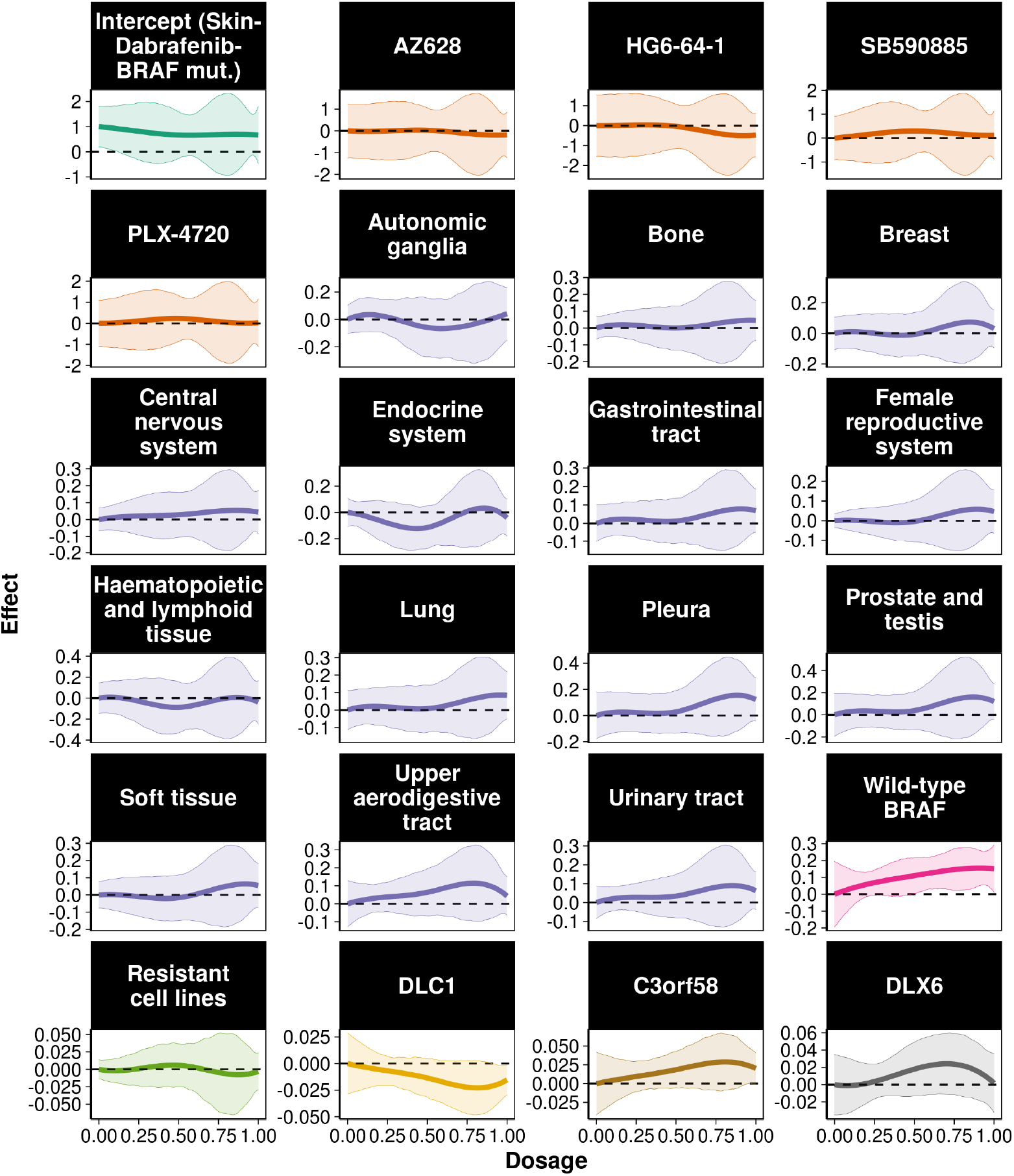
Estimated coefficient functions for the low-dimensional predictors and three of the selected genes. Estimated coefficient functions for intercept, different drugs, tissue of origin and three of the selected genes along with 95% bootstrap confidence intervals. Baseline corresponds to *BRAF* mutant cell lines treated with Dabrafenib in skin tumours.

Coefficient function estimates provide a lot of information about the dosage, cancer type and genetic effects on drug response. Fig 2 illustrates the estimated coefficient functions for different drugs, cancer types and three genes in relation to the model’s intercept, Dabrafenib response in *BRAF* mutant cell lines originating from the skin (melanoma). Except from HG6-64-1, all other BRAF inhibitors (AZ628, SB590885 and PLX4720) showed no addition effect compared to this intercept. Similar patterns can be observed for cancer cell lines coming from most of the tissues examined. This result indicates that the examined drugs may have similar or worse behaviour over the different dosages for most of the examined cancer types. Interestingly, we observed greater efficacy (negative values of the coefficient function) for cell lines originating from the endocrine system, autonomic ganglia and heamatopoietic and lymphoid tissues at lower dosages. The observed effect in endocrine system cell lines reflects the Dabrafenib responses observed in anaplastic thyroid cancer patients [32]. Interestingly, the drug, Trametinib, taken in combination with Dabrafenib is a MEK inhibitor, and genes interacting with MEK (*MAP2K1*) were selected features from our model (Fig 3). Together these results provide important insights into the effectiveness of the five *BRAF* targeted drugs examined on different cancer types, highlighting the potential for effective treatment of a wide range of cancers given the tumour genetic characteristics.

**Fig 3.**
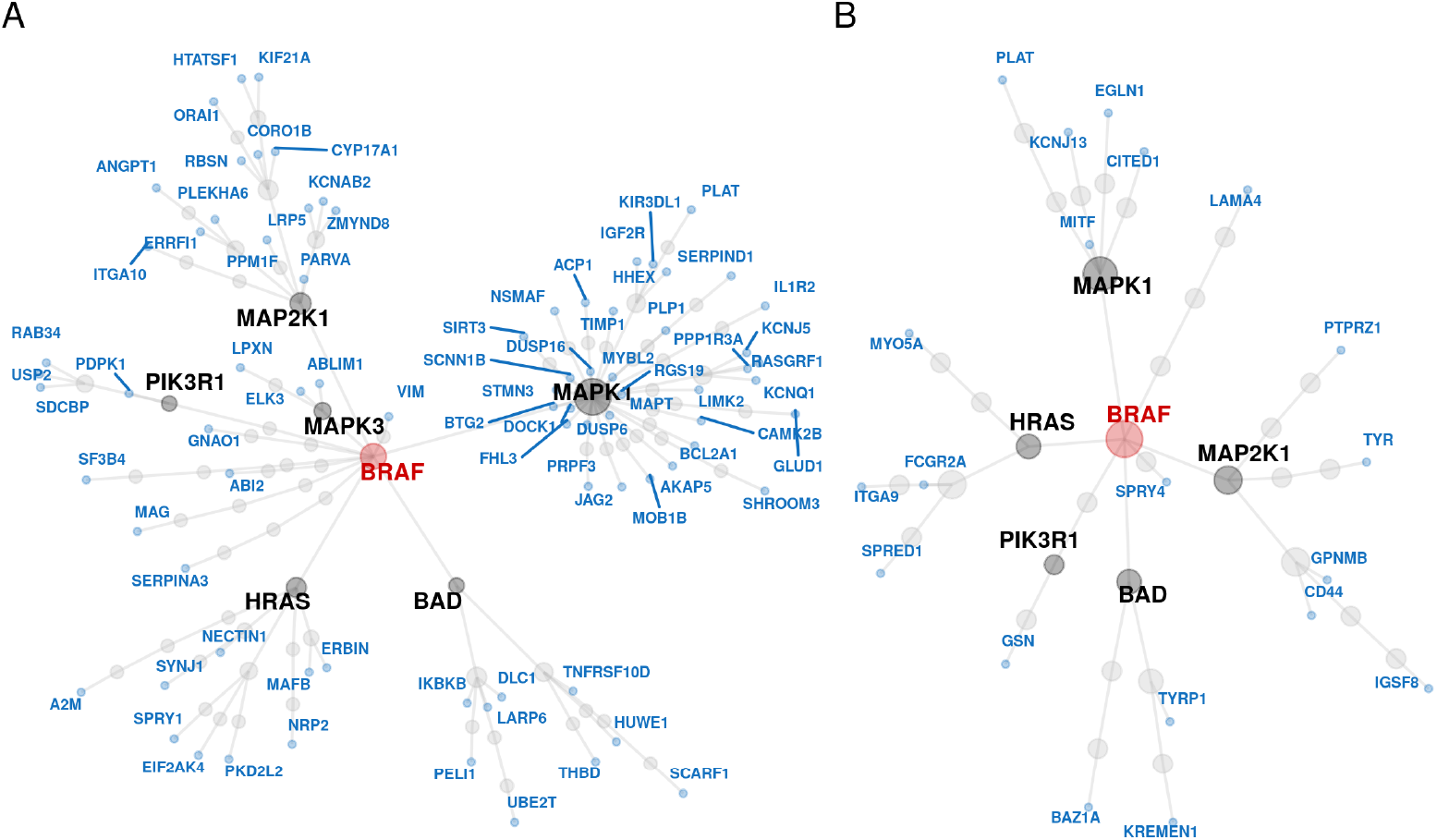
Protein-protein interaction network for the genes selected from the two-stage variable selection algorithm. (A) Undirected protein-protein interaction network between the 230 selected (blue) and the *BRAF* (red) genes (full scale analysis). (B) Undirected protein-protein interaction network between the 65 genes selected from the two-stage variable selection algorithm for the cell lines resistant to BRAF inhibitors (blue) and the *BRAF* (red) gene. In both panels genes depicted with black are the interaction mediators. Common mediators include the *HRAS, MAPK1, MAP2K1* and *BAD* genes.

Since the *BRAF* gene is the target of the drugs, mean fold change and protein-protein interaction network distance were used to examine whether and how the selected genes are related to inhibitors’ target. From the selected genes, 120 genes had a mean fold change greater than 1 whereas the rest had a mean fold change between 1 and 0.792. Some of the genes with the highest mean fold change of *BRAF* mutation were *PSMC3IP, KIF3C, UBE2Q2, SERPIND1* and *PLAT*, however only *PLAT* is displayed in Table 1. From the genes identified through the two-stage algorithm, 35% of them encode proteins interacting with the *BRAF* gene, though none of them directly. Most of the selected genes interact with the *BRAF* gene via pathways mediated by *HRAS, MAPK1* (*ERK*), *MAP2K1* (*MEK*) and *BAD* (Fig 3).

Since *HRAS* mutations are frequent in patients receiving *BRAF* targeted therapies [33], we examined the mean estimated trajectory over different dosages under treatment with *BRAF* inhibitors tested in six cancer cell lines with and without *BRAF* and *HRAS* mutations (Fig 4). As stated previously, we observed that in most cases HG6-64-1 seems to be the most effective drug. The estimated coefficient functions facilitate drug examination and response prediction under the different dosages. In some instances, we observed different drugs having similar behaviour for lower drug dosages and larger divergence for higher dosages. In most cases, regardless of the cell line origin, our method successfully estimates the expected survival rates of the cancer cell lines for the different drugs given their gene expression information.

**Fig 4.**
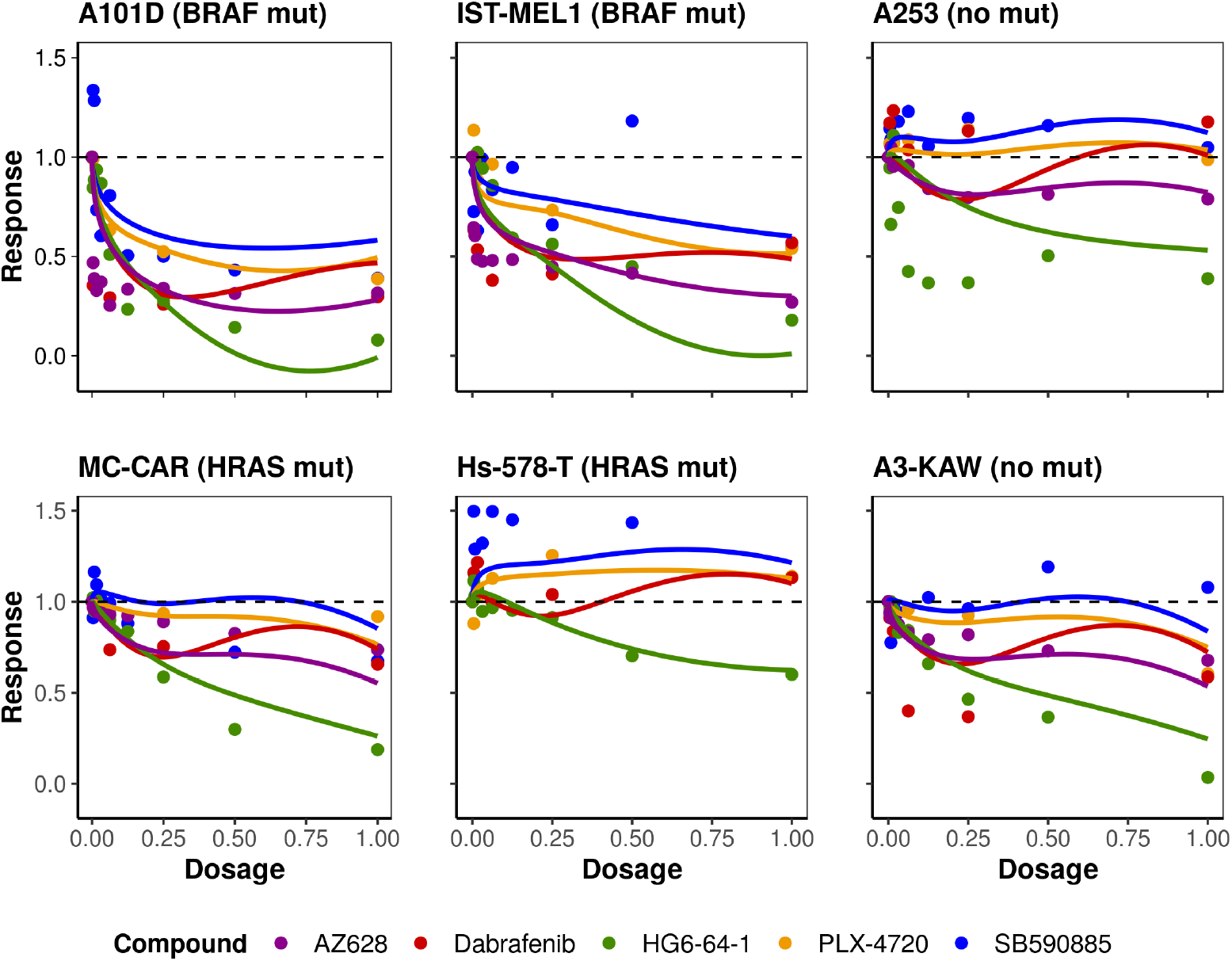
Estimated mean drug response trajectories for six cancer cell lines with *BRAF* and *HRAS* mutations. Observed responses (points) and estimated mean trajectory (lines) of cells’ concentration for cancer cell lines with and without *BRAF* and *HRAS* mutations after treatment with the five anticancer compounds examined.

For validation purposes, we performed the same analysis using the independently-generated GDSC2 dataset, with a different set of drugs. Note that the drug set in GDSC2 only partially overlaps with the one used in GDSC1. The results are reported in supplementary Fig S11, and show similar properties to the analysis of the BRAF drugs in Fig 4. As the GDSC2 dose-responses are produced from independently-generated experiments, the measured drug response is different for some of the drug-cell line combinations. We observe some divergences between GDSC1 and GDCS2 estimated trajectories for Dabrafenib and PLX-4720, which can be explained due to measurement error in the GDSC experiments themselves.

### Variable selection algorithm identifies cancer pathways associated with BRAF inhibitor response

Using our functional regression approach, we identified 230 genes that were selected via the SCAD step (observed gene set). We used the Enrichr [34,35], WikiPathways [36] databases to see if the selected genes can be grouped into common functional classes or pathways. In total, 183 pathways identified, from which 11 were statistically significant at 5% level, including apoptosis modulation, NOTCH1 regulation, and MAPK signaling (Table S6). The model identified genes (*IKBKB, RASGRF1, DUSP16, DUSP8, DUSP6, MAPT* and *IL1R2*) downstream of the MAPK signaling pathway targeted by BRAF inhibitors.

Previous studies of these pathways have found associations with tumourgenesis and cancer treatment [37–40]. Genes in more than one of these pathways include *IKBKB, PLAT, IL1R2* and *PDPK1*. The IKB kinase composed of IKBKB had previously been suggested as a marker of sensitivity for combination therapy with BRAF inhibitors [41]. Taken together, these results suggest that the identified associations between the drug response and the observed genes may reveal new predictive markers of tumour response to the examined BRAF inhibitors.

In addition to the pathway enrichment analysis, we used the Molecular Signatures Database (MSigDB database v7.0 updated August 2019: [42]) to compute overlaps between the observed gene set and known oncogenic gene sets. Fig 5 displays the 29 overlaps found. Interestingly, we identified three instances where the observed gene set significantly overlapped with gene sets over-expressing an oncogenic form of the *KRAS* gene.

**Fig 5.**
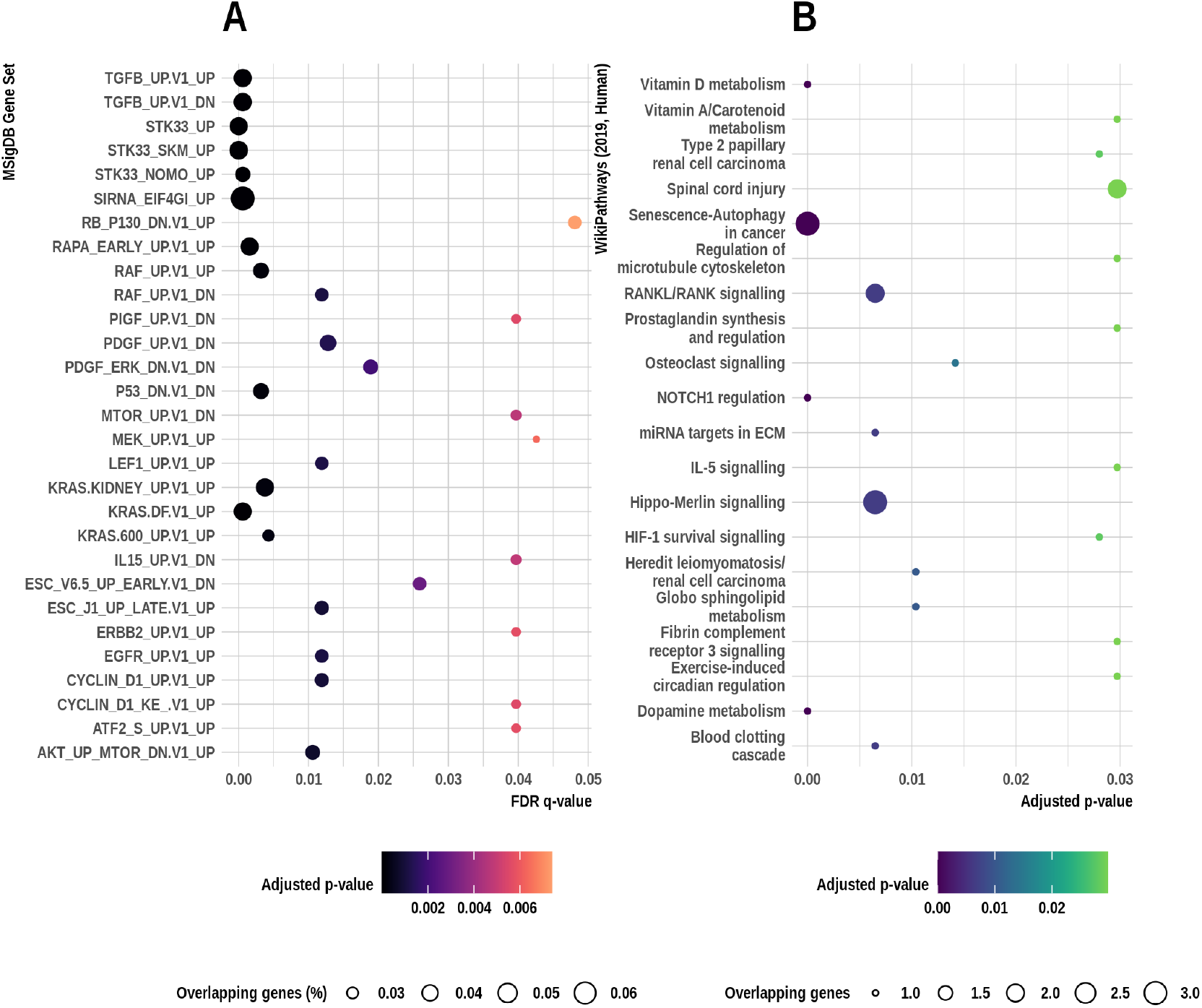
Overlaps between the observed gene set and oncogenic signatures in the Molecular Signatures Database (full data analysis); signalling pathways enriched for genes predictive of *BRAF* inhibitor response (resistant cell lines). (A) Full gene set names can be found in Table S8. Overlaps have been detected using gene set enrichment analysis performed using a hypergeometric distribution. The false discovery rate analog of the hypergeometric p-value is displayed after correction for multiple hypothesis testing according to Benjamini and Hochberg [43]. (B) Top 20 enriched signalling pathways along with the adjusted p-values and the number of overlapping genes obtained after pathway enrichment analysis to the resistant cell line analysis results (for full list of the pathways identified see Table S7).

We further explored potential biologically relevant pathways using the Reactome database [44,45]. More than 40 enriched pathways were identified at a 5% significance level. The top 40 pathways are depicted in Fig 6 along with the pathway-gene network of the top 5 pathways (for the full list, see Table S12). Interestingly, axon guidance and VEGF signalling were among the enriched pathways, confirming relevance of the selected genes to the intended role of the examined compounds, since BRAF kinase activity drives axon growth in the central nervous system [46] and VEGF blockade has potential anti-tumour effects when combined with BRAF inhibitors [47]. Note that axon growth has not thus far been directly implicated in BRAF inhibitor activity, and thus our analysis provides important evidence towards this theory.

**Fig 6.**
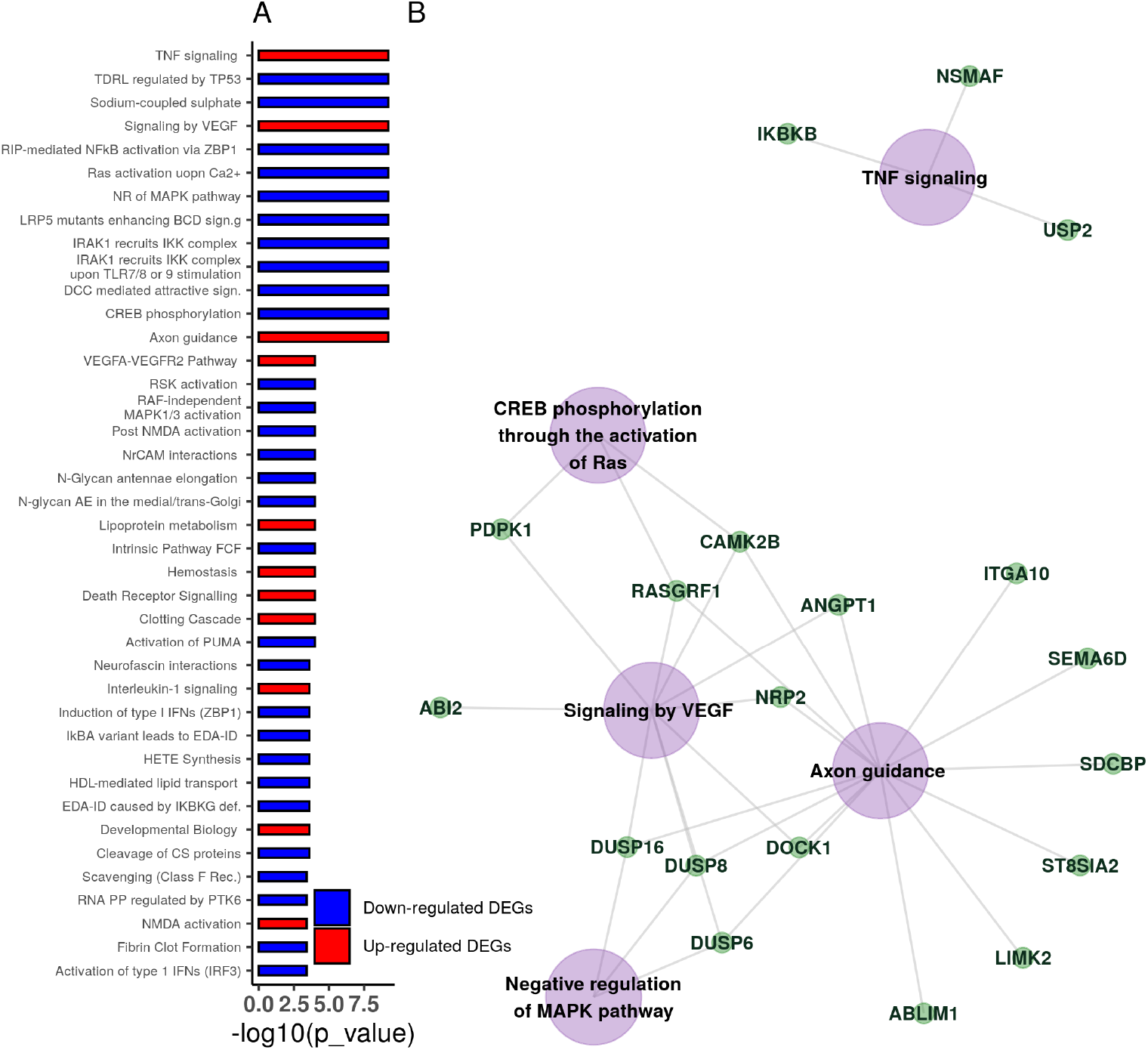
Pathway enrichment analysis using the Reactome database. (A) Top 40 enriched signalling pathways along with the adjusted p-values. For the full list, see Table S12. (B) Pathway-gene network of the top 5 enriched signalling pathways as found using Reactome [44, 45].

### Identifying dose-dependent genes in drug-resistance conditions

Acquired resistance to BRAF inhibitors is often observed in the clinic [48]. To further examine the utility of the employed methodology, we applied the variable selection algorithm to a data subset containing only cell lines with mutations activating resistant mechanisms to BRAF inhibitors [49]. Out of the 951 cell lines in the data, 191 had some mutation in any of the following: *RAC1* gene, *NRAS* gene, *cnaPANCAN44* or *cnaPANCAN315*. We identified 65 genes associated with dose-response, though none of them were directly associated with the MAPK/ERK pathway. However, from these, 25 genes have been found to indirectly interact with the *BRAF* gene (Fig 3) and 21 to overlap with three oncogenic gene sets in the Molecular Signatures Database (genes down-regulated in NCI-60 panel of cell lines with mutated *TP53*; genes up-regulated in Sez-4 cells (T lymphocyte) that were first starved of *IL2* and then stimulated with *IL21*, and; genes down-regulated in mouse fibroblasts over-expressing *E2F1* gene; Table S9). Finally, we found 34 pathways enriched for genes predicting drug response of the mutated cell lines to the examined BRAF inhibitors, of which the top 20 are depicted in Fig 5(B).

Table 2 presents gene rankings based on the AUC and the overall coefficient function effect (sign) for the 42 genes in either the enriched pathways, the three oncogenic gene sets discussed above or the protein-protein interaction network with the *BRAF* gene (full list available in Table S5). Eight of the selected genes in the current implementation were also selected from the algorithm implemented on the full data: *ASB9, PRSS33, GJA3, PLAT, KLF9, BFSP1, MTARC1* and *UCN2*.

**Table 2.**
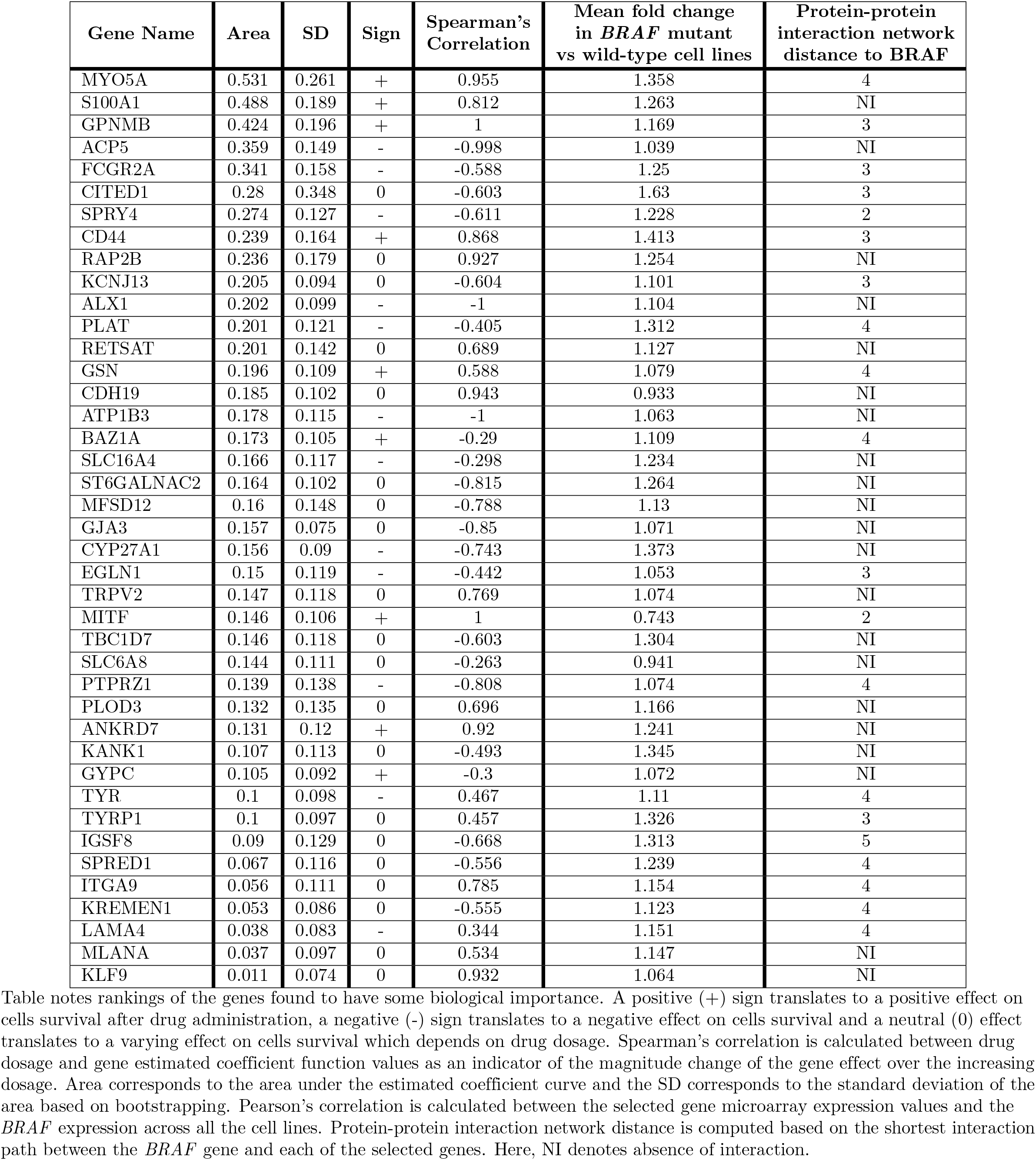
Rankings of the genes identified from the pathway and oncogenic gene set enrichment analysis.

### Predictive performance of dose-dependent models

As discussed above, the employed methodology gives a good overview of the baseline genetic effect on drug response. We assessed the overall predictive performance of our method using 10-fold cross validation under two different scenarios. For the first, we split the data into training and test set holding out the experimental units (cancer cell line-drug combinations) and for the second, holding out cancer cell lines. The absolute mean error for both cases was around 0.12. Our analysis shows robust cross-validated performance when it comes to predicting sensitivity to the administered drugs, as shown in Fig S3 which displays the correlation between predicted and true response. This result was further validated by repeating the analysis on the independently-generated GDSC2 dataset, using a different set of drugs (Fig S10), which demonstrated comparable predictive performance.

Predictive accuracy for the dose-response curves was evaluated under four different sub-scenarios: prediction of the most effective drug-dosage combination for the 951 cell lines in the data set; prediction of the most effective drug given a cell line; prediction of the most effective dosage given treatment with a particular drug and prediction of the most effective dosage range given treatment with a drug (Table 3). The proposed model performs well when it comes to predicting the most effective drug or dosage range (≈79% in both scenarios). Results are less reliable when it comes to prediction of the exact dosage or drug-dosage combination (≈48-49% and ≈57-58% in both scenarios) but this can be due to either the large variability observed in the observed responses or due to the small number of cell lines for some predictor level combinations. Results were similar for both cross-validation scenarios (differences range from 0 to <2%, Table 3), meaning that as long as a cell line has similar genetic characteristics to those observed, the model can be reliable in predicting the outcome after anticancer drug administration.

**Table 3.**
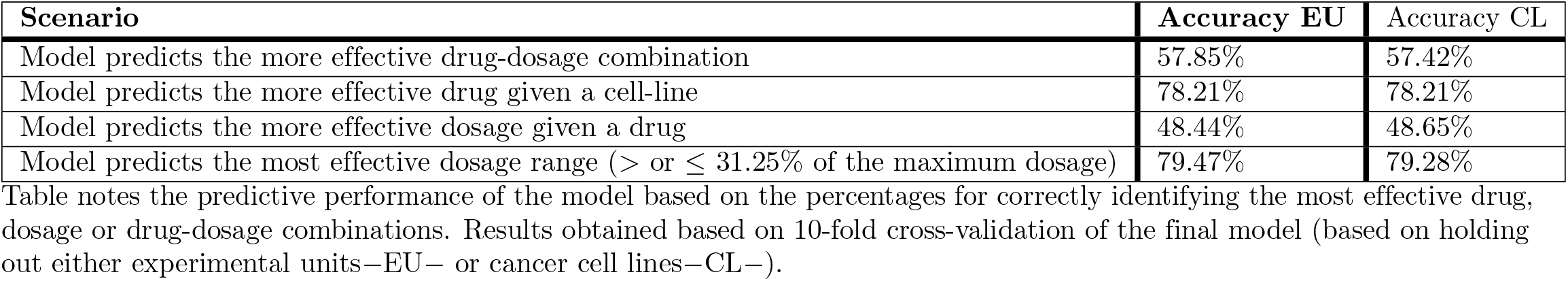
Predictive performance of the employed model (mean absolute error=0.121).

We additionally compared the performance of our two-stage algorithm approach to a penalized linear (LASSO) regression for predicting the IC50 and area under the dose-response curve (AUC). Note that our functional regression model is not directly predicting either of these values, but rather predicts the full drug-response curve. As this is a harder problem, we would expect the LASSO to have a natural advantage; however, our method has the added benefit of being able to detect dose-dependent associations, which is not possible when predicting summary statistics of the dose-response curve directly. We employed 10-fold cross-validation to evaluate the predictive error in terms of root mean squared error (RMSE), and we used a sigmoid curve fit for estimating the IC50 values from the predicted dose-response curves with our two-stage method. Our method outperformed standard LASSO in terms of predicting the AUC (RMSE_2–*stage*_ = 0.176; RMSE*_LASSO_* = 0.347) and performed well on predicting the IC50, although the LASSO performed better (RMSE_2–*stage*_ = 1.969; RMSE*_LASSO_* = 1.134). This could be expected, as estimating the IC50 from the predicted dose-response curve adds a further level of complexity, compared to directly predicting this value using the LASSO.

### Conclusion

Genetic alternations and gene expression in tumours are known to affect disease progression and response to treatment. Here, we studied dosage-dependent associations between gene expression and drug response, using a functional regression approach which adjusts for genetic factors. We analysed data from the Genomics of Drug Sensitivity in Cancer project relating to drug effectiveness for suspending cancer cell proliferation under different dosages, and examined five *BRAF* targeted inhibitors, each applied in a number of common and rare types of cancer cell lines. Our implementation of a two-stage screening algorithm revealed a number of genes that are potentially associated with drug response. Gene, drug and cancer type trajectories have been modelled using a varying coefficient modelling framework. The proposed methodology allows for dose-dependent analysis of genetic associations with drug response data. It enables us to study the effect of different drugs simultaneously, which results in high accuracy of drug response prediction. Drug comparisons using the proposed methodology could support drug repositioning, especially in disease indications where existing treatment options are limited. In addition, our methodology can help to reveal unknown potential relationships between genetic characteristics and drug efficacy. Hence, the good predictive performance of our method could be due to the fact that some genes may act as proxies for unmeasured phenotypes that are directly relevant to drug sensitivity.

Our work relies on two major assumptions. First, that out of tens of thousands genes regulating protein composition only a small proportion is actually associated with cancer cell survival in a dosage-dependent manner. In other words, transcriptomic profiles exert influence on disease progress after drug administration in a sparse and dynamic way. However, if a large number of genes are associated with the drug response, our method may produce biased results, and some important information about the biological mechanisms can be lost. Secondly, we assume that the different drugs are comparable on the scale of maximum dosage percentage level for our joint model. However, we acknowledge that different drugs have different chemical structure and maximum screening concentrations. Our focus is to identify genetic components that could be informative for dose response given drugs that belong to a particular family, for example *BRAF* targeted therapies. However, our methodology is flexible enough to allow each drug to be examined separately if it appears to be clinically appropriate.

Drug response prediction from gene expression data has been widely studied in the literature. Sparse regression methods, gene selection algorithms such as the Ping-pong algorithm [50], or a combination of network analysis and penalised regression, e.g. the sparse network-regularized partial least squares method [51], have all been employed to simultaneously predict drug response and select genetic factors that seem to be associated with the drug response. However, none of these methods are able to quantify the effect of drug dosage on the response, which is one of the main contributions of this work. Employing the proposed dose-varying model gives a detailed picture of different drug effects and can be extremely valuable in predicting drug response for agents with small therapeutic range and high toxicity levels. Our algorithm showed moderate predictive performance due to the complexity of predicting whole drug-response curves. Methods for further enhancing the performance of the proposed methodology, such as judicious use of prior information and leveraging information sharing across multiple data sources should be explored in the future in order to overcome this issue and make good use of its full potentials.

To conclude, the main purpose of this paper is to examine the dose-dependent associations between genes and drugs. The proposed methodology, by using the raw data to infer the effects of interest, allows to obtain a more comprehensive picture of the biological mechanisms that undergo cancer treatment and the role of drug dose on that. In addition, due to its simple structure, it allows extension to different types of molecular data (e.g. RNA-seq gene expression, methylation or mutational profiles) and enrichment with further information, such as drug chemical composition.

## Supporting information captions

**S1 Text. Accurate detection of drug associated genes from simulated responses.** Simulated responses have been generated to examine the accuracy of the employed method in detecting the genes that are truly associated to drug response. Three screening thresholds, three active gene sets and two covariance structure scenarios for the repeated measurements simulation have been considered. This text includes all the details of the simulation study that we conducted.

**S2 Fig. Distribution of tissue of origin across the five BRAF compounds used for cell line screening in the Genomics of Drug Sensitivity in Cancer data.** Overall, similar proportion of cell lines have been treated with all of the compounds examined with smaller number of cell lines been treated with AZ628, Dabrafenib and PLX-4720. Larger number of cell lines in the data set were originated from the lungs, the gastrointestinal tract and the haematopoietic and lymphoid tissues.

**S3 Fig. Prediction accuracy for each different drug and scenario.** Pearson correlation was estimated across observed and predicted AUC values. AUC values have been computed by calculating the area under the coefficient function curve (both observed and predicted). Training and test sets have been considered based on either the experimental units or on cancer cell lines only.

**S4 Table. Full gene rankings based on the estimated area under the coefficient function curve (analysis on the full data set).** Gene rankings of all selected genes based on the magnitude of the genetic effect on drug response. A positive (+) sign translates to a positive effect on cells survival after drug administration, a negative (-) sign translates to a negative effect on cells survival and a mixed (0) effect translates to a varying effect on cells survival which depends on drug dosage.

Spearman’s correlation is calculated between drug dosage and gene estimated coefficient function values as an indicator of the magnitude change of the gene effect over the increasing dosage. Area corresponds to the area under the estimated coefficient curve and the SD corresponds to the standard deviation of the area based on bootstrapping. Mean fold change is calculated between the selected gene expression values of the cell lines carrying BRAF mutations with respect to wild type. Protein-protein interaction network distance is computed based on the shortest interaction path between the BRAF gene and each of the selected genes. Here, NI denotes absence of any interaction.

**S5 Table. Full gene rankings based on the estimated area under the coefficient function curve (analysis on resistant cell lines).** Gene rankings of all selected genes based on the magnitude of the genetic effect on drug response. A positive (+) sign translates to a positive effect on cells survival after drug administration, a negative (-) sign translates to a negative effect on cells survival and a mixed (0) effect translates to a varying effect on cells survival which depends on drug dosage.

Spearman’s correlation is calculated between drug dosage and gene estimated coefficient function values as an indicator of the magnitude change of the gene effect over the increasing dosage. Area corresponds to the area under the estimated coefficient curve and the SD corresponds to the standard deviation of the area based on bootstrapping. Mean fold change is calculated between the selected gene expression values of the cell lines carrying *BRAF* mutations with respect to wild type. Protein-protein interaction network distance is computed based on the shortest interaction path between the BRAF gene and each of the selected genes. Here, NI denotes absence of any interaction.

**S6 Table. Signalling pathways linked to genes predictive of BRAF inhibitor response (analysis on the full data set).** Signalling pathways along with the adjusted p-values and the number of overlapping genes obtained after pathway enrichment analysis to the full scale analysis results.

**S7 Table. Signalling pathways linked to genes predictive of BRAF inhibitor response (analysis on resistant cell lines).** Signalling pathways along with the adjusted p-values and the number of overlapping genes obtained after pathway enrichment analysis to the resistant cell line analysis results.

**S8 Table. Overlaps between the observed gene set and oncogenic signatures in the Molecular Signatures Database (analysis on the full data set).** Overlaps have been detected using gene set enrichment analysis performed using a hypergeometric distribution. The false discovery rate analog of the hypergeometric p-value is displayed after correction for multiple hypothesis testing according to Benjamini and Hochberg.

**S9 Table. Overlaps between the observed gene set and oncogenic signatures in the Molecular Signatures Database (resistant cell lines analysis).** Overlaps have been detected using gene set enrichment analysis performed using a hypergeometric distribution. The false discovery rate analog of the hypergeometric p-value is displayed after correction for multiple hypothesis testing according to Benjamini and Hochberg.

**S10 Fig. Prediction accuracy for each different drug and scenario: analysis performed on GDSC2 data.** Pearson correlation across observed and predicted AUC values. AUC values have been computed by calculating the area under the coefficient function curve (both observed and predicted) using the GDSC2 data. Training and test sets have been considered based on either the experimental units or on cancer cell lines only.

**S11 Fig. Estimated mean drug response trajectories for *BRAF* and *HRAS* mutated and non-mutated cancer cell lines: analysis performed on GDSC2 data.** Observed responses (points) and estimated mean trajectory (lines) of cells’ concentration for cancer cell lines with and without *BRAF* and *HRAS* mutations after treatment with the eight anticancer compounds examined using data from GDSC2.

**S12 Table. Signalling pathways linked to genes predictive of BRAF inhibitor response (analysis on the full data set)-Reactome.** Signalling pathways along with the adjusted p-values and the number of overlapping genes obtained after pathway enrichment analysis to the full scale analysis results using the Reactome database.

## Acknowledgments

We acknowledge Emily Chambers and the Sheffield Bioinformatics Core Facility for their assistance in GDSC2 data pre-processing. Their input to this study is greatly appreciated.

