## Supplementary figures and images for "A regularized functional regression model enabling transcriptome-wide dosage-dependent association study of cancer drug response"

### Supporting Information Figure S2

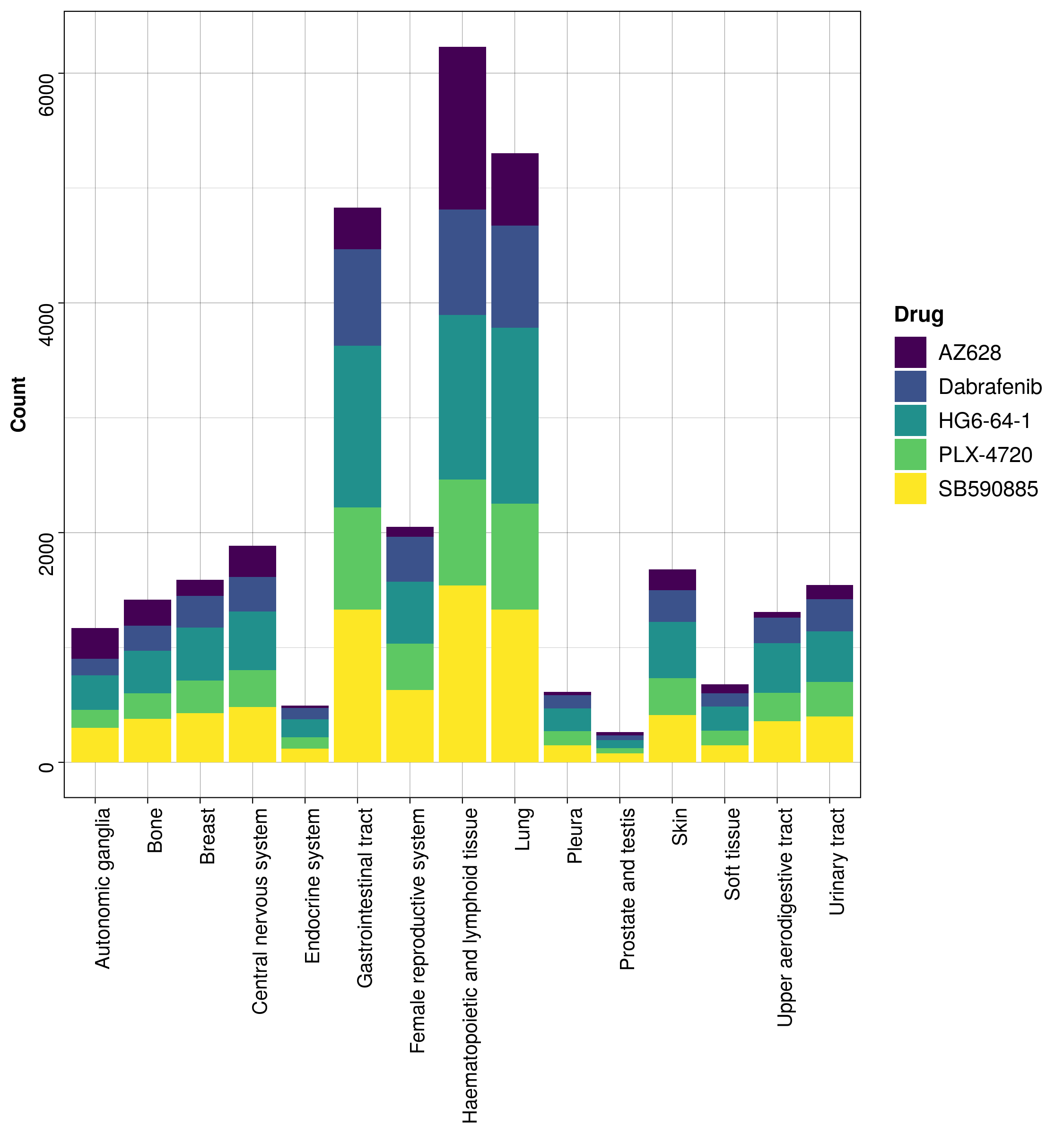

### Supporting Information Figure S3

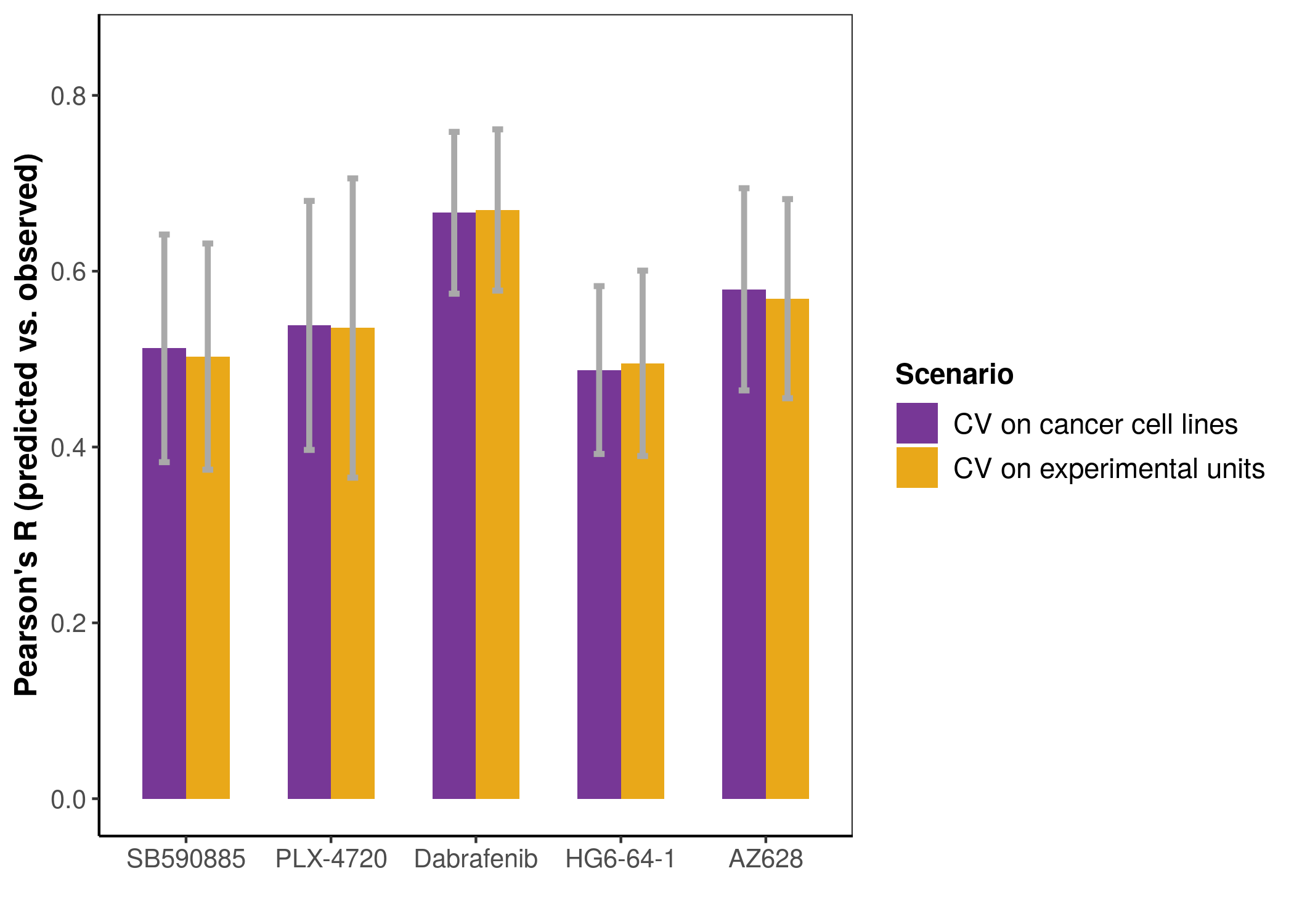

### Supporting Information Figure S10

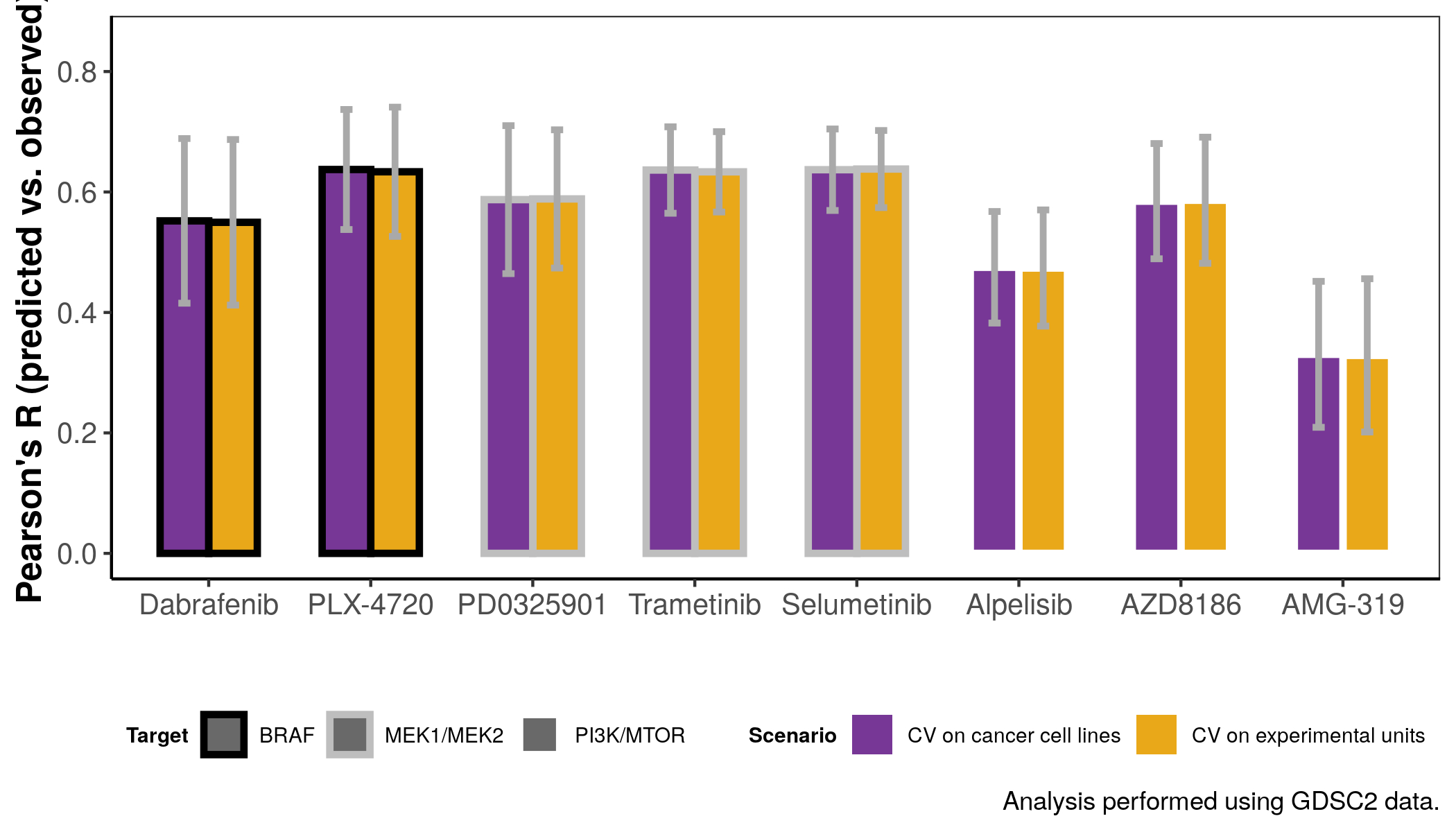

### Supporting Information Figure S11

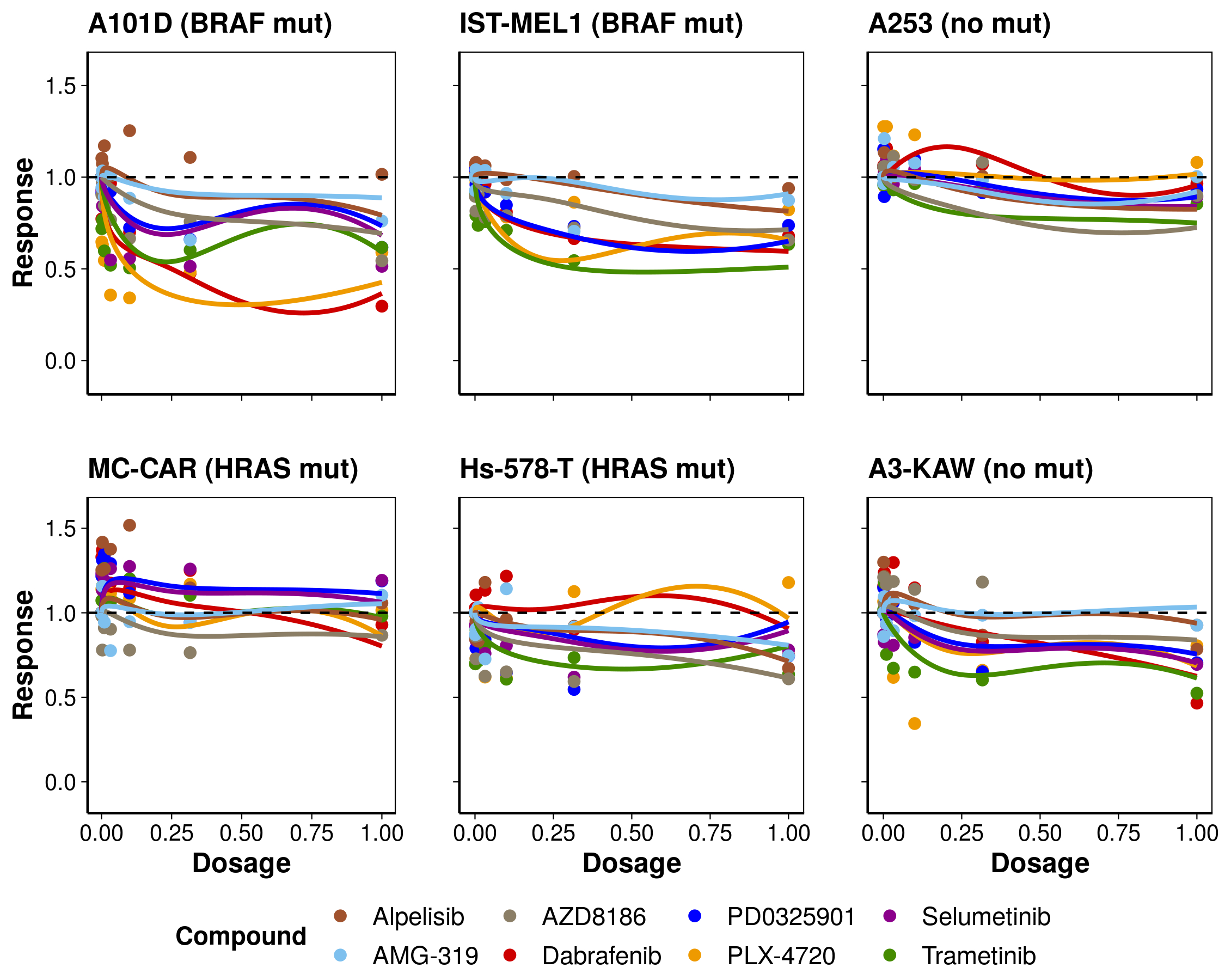
