## Supporting Information Text S1 for "A regularized functional regression model enabling transcriptome-wide dosage-dependent association study of cancer drug response"

Biomarker detection for revealing anticancer drug dynamics

Evanthia Koukouli[1*], Dennis Wang [2,3], Frank Dondelinger[4◑], Juhyun Park[1◑]

**1** Department of Mathematics and Statistics, Fylde College, Lancaster University, Bailrigg, Lancaster, UK

**2** Sheffield Institute for Translational Neuroscience, University of Sheffield, Sheffield, UK

**3** Department of Computer Science, University of Sheffield, Sheffield, UK

**4** Centre for Health Informatics and Statistics, Lancaster Medical School, Lancaster University, Bailrigg, Lancaster, UK

### Introduction

Cancer is a heterogeneous disease, with individual tumours showing sometimes very different mutational and molecular profiles. The genetic makeup of a tumour influences how it reacts to a given anti-cancer drug. However, due to lack of predictive markers of tumour response, often patients with very different tumour genetic makeup will receive the same therapy, resulting in high rates of treatment failure [1]. Large clinical trials in rapidly lethal diseases are expensive, complex and often lead to failure due to lack of efficacy [2]. Therefore, there is a need for more effective and personalised therapeutic strategies that can improve cancer treatment decisions, and hence patient outcomes.

One major issue for some cancer treatments, e.g. chemotherapies, are cytotoxic effects that result in collateral damage of the healthy host tissue [3]. Patient remission depends not only on the selection of the best therapeutic agent but also on the determination of the optimal dosage, especially when drugs with small therapeutic range, high toxicity levels or both are administered. Genetic factors can help fine-tune the dosage for individual patients, so that the minimal effective dosage can be delivered [4]. Previous work has examined the difference in transcriptional response [5] and drug response at the cell population level after administering anticancer drugs in various dosages [6, 7].

Cancer cell line drug screens provide valuable information about biomarkers that are predictive of drug response. During the last decade, there have been several systematic studies aiming to examine pharmacogenomic relationships [8–11]. These studies were conducted on human cancer cells that have been isolated from affected tissues, grown in vitro and treated with anti-cancer inhibitors. By examining the genomic profiles of these cell lines, investigators were able to identify relationships between cancer-driven genetic alterations and drug response. However, these relationships have only been modelled on the aggregate response, and hence little is known about the relationship between drug dosage and genetic factors. Recently, Tansey et al. [12] proposed a method for modelling drug-response curves via Gaussian processes and linking them to biomarkers using a neural network prediction model. The authors did not use their model for dosage-dependent inference of biomarker effects, and the highly non-linear neural network model makes interpretation of biomarker effects challenging.

Gene expression profiles can provide valuable functional information on the genetic mechanisms which determine anti-cancer drug response, offering more tailored treatments where common therapies become ineffective. However, statistical analysis for linking transcriptomic profiles with drug response becomes challenging due to the high-dimensional nature of the data. Over the last 20 years, researchers developed statistical methodologies not only to mitigate the problem of high dimensionality [13–18], but also to detect markers of positive drug response to cancer treatment [19,20] and predict patient response after drug administration [1,5,16,21–25]. While these previous methods have gone some way towards solving the challenges associated with drug response modelling, none of them address all of the issues that arise in personalised medicine, namely: selecting genes associated with drug response, identifying the optimal dosage, characterising gene-dose relationships and predicting response for one or multiple drugs.

With regards to the high-dimensional nature of the dataset, it is worth noting that highly-complex data sets with non-stationary trends are not easily amenable to analysis by classic parametric or semi-parametric mixed models. However, the effect of genes on drug response over different drug dosages (dose-varying effect) can be examined using varying coefficient models which allow for the covariate effect to be varying instead of constant [26]. Methods to estimate the covariate (e.g. gene) effect include global and local smoothing e.g. kernel estimators [27,28], basis approximation [29] or penalised splines [30]. The most straightforward and computationally efficient method is through basis approximation where each coefficient function is approximated through some basis functions and the varying coefficient model can be written as a linear regression model. Then, estimation for repeated measurements data (e.g. drug response over different dosages) can be incorporated through minimising a weighted least squares criterion based on a specified weighting scheme (repeated measurements covariance structure) [29]. However, inference becomes impossible as the number of predictors increases and when selecting a smaller number of important variables for inclusion into the model is clinically beneficial. Sparse regression has enabled a more flexible and computationally "inexpensive" way of choosing the best subset of predictors. When combining sparse regression with the varying coefficient model framework, predictors are handled jointly under the assumption that the majority are irrelevant to the outcome variable. Penalties from group versions of the least absolute shrinkage and selection operator (LASSO), smoothly clipped absolute deviation (SCAD), bridge etc. have been used for fitting the varying coefficient model [31]. Because these methods handle all of the predictors jointly, their implementation becomes extremely challenging and impractical when the number of predictors (e.g. thousands) is much larger than the number of samples (e.g. hundreds). Consequently, attempts to develop prior univariate tests focused on filtering out the unimportant predictors by simply estimating the association of each predictor to the outcome variable separately [32–34]. Often, these screening methods are conservative, and still return many more predictors than those which are truly associated to the response. To overcome this issue, regularisation or alternative variable selection methods have been used after screening to further fine-tune the set of predictors [32, 34].

The advantage of using varying coefficient models along with a variable screening algorithm on genomic data sets was first introduced to explore the effect of genetic mutations on lung function [32]. Here, we extended their methodology to a completely different objective of assessing the transcriptomic effect on anti-cancer drug response, where our coefficient functions were allowed to vary with dosage. Note that unlike in Tansey et al. [12], biomarker effects will be a linear function function of dosage, allowing for straight-forward interpretation of the coefficient functions.

We developed a functional regression framework to study the effectiveness of multiple anticancer agents applied in different cancer cell lines under different dosage levels, adjusting for the transcriptomic profiles of the cell lines under treatment. We considered a dose-varying coefficient model, along with a two-stage variable selection method in order to detect and evaluate drug-gene relationships. We applied this method to data extracted from the Genomics for Drug Sensitivity in Cancer (GDSC) project [9]. To compare and differentiate similar treatments, we examined the effect of five BRAF targeted compounds under different dosages to almost 1000 cancer cell lines. We used baseline gene expression measurements for the cancer cell lines to investigate gene-drug response relationships for almost 18000 genes. Gene rankings were obtained based on the gene effect on the drug response. Consequently, in contrast to past studies, we managed to model the whole dose-response curve, rather than a summary statistic of drug response (e.g. IC50), which allowed us to identify trends in the gene-drug association at untested dose concentrations.

$i = 1, \ldots, 3805, j = 1, \ldots, n_i$.

### A two-stage algorithm for identification of gene-drug associations

Let the repeated measures data $\{(d_{ij},\ y_{ij},\ \boldsymbol{z}_i,\ \boldsymbol{x}_i) : j = 1, \ldots, n_i,\ i = 1, \ldots, n\}$, where $y_{ij}$ is the response of the $i$th experimental unit (corresponds to a drug sensitivity assay of a specific drug on a specific cell line) at the $j$th drug dosage level $d_{ij}$ and $\boldsymbol{z}_i$ along with $\boldsymbol{x}_i$ are the corresponding vectors of scalar (dose-invariant) covariates. The covariate vector $\boldsymbol{z}_i = (1, z_{i1}, \ldots, z_{ip})^T$ is a low-dimensional vector of predictors that should be included in the model, whereas $\boldsymbol{x}_i = (x_{i1}, x_{i2}, \ldots, z_{iG})^T$ is a high-dimensional vector, i.e. 17737 gene expression measurements, that needs to be screened. We assumed that only a small number of $x$-variables (in our case, genes) are truly associated with the response while most of them are expected to be irrelevant; i.e. we make a sparsity assumption.

To explore potential dose-varying effects between the covariates and the drug response, we consider the following varying coefficient model:

$$y_{ij} = \sum_{k=0}^{p} z_{ik}\beta_k(d_{ij}) + \sum_{g=1}^{G} x_{ig}\gamma_g(d_{ij}) + \varepsilon_{ij} \tag{2}$$

where $\{\beta_k(\cdot), k = 0, \ldots, p\}$ and $\{\gamma_g(\cdot), g = 1, \ldots, G\}$ are smooth functions of dosage level $d \in \mathscr{D}$, where $\mathscr{D}$ is a closed and bounded interval of $\mathbb{R}$. The errors $\varepsilon_{ij}$ were assumed to be independent across subjects and potentially dependent within the same subject with conditional mean equal to zero and variance $\mathrm{Var}(\varepsilon) = \sigma^2(d) = V(d)$.

Methods for estimating the coefficient functions in Eq (2) include local and global smoothing methods, such as kernel smoothing, local polynomial smoothing, basis approximation smoothing etc. Due to computational convenience, for this application we used basis approximation smoothing via B-splines.

Let the sets of basis functions $\{B_{lk}(\cdot) : l = 1, \ldots, L_k\}$ and $\{B'_{lg}(\cdot) : l = 1, \ldots, L_g\}$ and constants $\{\zeta_{lk} : l = 1, \ldots, L_k\}$ and $\{\eta_{lg} : l = 1, \ldots, L_g\}$ where $k = 0, \ldots, p$ and $g = 1, \ldots, G$ such that, $\forall d \in \mathscr{D}$, $\beta_k(d)$ and $\gamma_g(d)$ can be approximated by the expansion

$$\beta_k(\cdot) \approx \sum_{l=1}^{L_k} \zeta_{lk} B_{lk}(\cdot) \ \text{ for } \ k = 0, \ldots, p \tag{3}$$

$$\gamma_g(\cdot) \approx \sum_{l=1}^{L_g} \eta_{lg} B'_{lg}(\cdot) \text{ for } g = 1, \ldots, G. \tag{4}$$

Substituting $\beta_k(\cdot)$ and $\gamma_g(\cdot)$ of Eq (2) with Eq (3) and Eq (4), we approximated Eq (2) by

$$y_{ij} \approx \sum_{k=0}^{p} z_{ik} \sum_{l=1}^{L_k} \zeta_{lk} B_{lk}(d_{ij}) + \sum_{g=1}^{G} x_{ig} \sum_{l=1}^{L_g} \eta_{lg} B'_{lg}(d_{ij}) + \varepsilon_{ij} \tag{5}$$

If $B_k(\cdot)$ and $B'_g(\cdot)$ are groups of B-spline basis functions of degree $q_k$ and $q_g$ respectively, and $\delta_0 < \delta_1 < \ldots < \delta_{K_k} < \delta_{K_k+1}$ and $\delta_0 < \delta_1 < \ldots < \delta_{K_g} < \delta_{K_g+1}$ are the corresponding knots, then $L_k = K_k + q_k$ and $L_g = K_g + q_g$.

Using the approximation Eq (5), the coefficients $\boldsymbol{\zeta} = (\zeta_0, \zeta_1, \ldots, \zeta_p)^T$ and $\boldsymbol{\eta} = (\eta_1, \eta_2, \ldots, \eta_G)^T$ can be estimated by minimizing the squared error

$$\ell_w((\boldsymbol{\zeta}, \boldsymbol{\eta})^T) = \sum_{i=1}^{n} \sum_{j=1}^{n_i} w_{ij} \left[ y_{ij} - \sum_{k=0}^{p} z_{ik} \sum_{l=1}^{K_k} \zeta_{lk} B_{lk}(d_{ij}) - \sum_{g=1}^{G} x_{ig} \sum_{l=1}^{L_g} \eta_{lg} B'_{lg}(d_{ij}) \right] \tag{6}$$

where $w_{ij}$ are known non-negative weights.

In cases where $p + G >> n$ though, minimisation of Eq (6) is infeasible. Our aim was to identify factors of the covariate vector $\mathbf{x} = (\mathbf{x}_1, \mathbf{x}_2, \ldots, \mathbf{x}_G)^T$ (genes) that are truly associated with the response (cancer cell line sensitivity to the drug). In addition, we wanted to explore potential dose-varying effects on the drug response.

We make the following sparsity assumption: any valid solution $\hat{\boldsymbol{\gamma}}(d)$ will have $\hat{\gamma_g}(d) = 0, \forall d \in \mathscr{D}$ for the majority of components $g$. To detect non-zero coefficient functions we applied a two-stage approach which incorporated a variable screening step and a further variable selection step.

### Screening

The sparsity assumption applies only to components of $\mathbf{x}$, the high-dimensional covariate vector in Eq (2).

Let the set of indices

$$\mathscr{M}_0 = \{1 \le g \le G : ||\gamma_g(\cdot)||_2 > 0\} \tag{7}$$

where $|| \cdot ||_2$ is the $L_2$-norm. In order to rank the different components of $\boldsymbol{x}$, we fitted the marginal non-parametric regression model for the $g$th $x$-predictor:

$$y_{ij} \approx \sum_{k=0}^{p} z_{ik} \sum_{l=1}^{K_k} \zeta_{lk}^{(g)} B_{lk}^{(g)}(d_{ij}) + x_{ig} \sum_{l=1}^{L_g} \eta_{lg}^{(g)} B_{lg}^{(g)\prime}(d_{ij}) + \varepsilon_{ij}^{(g)} \tag{8}$$

where: $\{B_{lk}^{(g)}(\cdot) : l = 1, \dots, L_k\}$ and $\{B_{lg}^{(g)\prime}(\cdot) : l = 1, \dots, L_g\}$ are sets of coefficient functions; $\{\zeta_{lk}^{(g)} : l = 1, \dots, L_k\}$ and $\{\eta_{lg}^{(g)} : l = 1, \dots, L_g\}$ are constants to be estimated, $k = 0, \dots, p$; and, $\varepsilon^{(g)}$ is the error term similar to Eq (5). We then computed the following weighted mean squared error for each $g \in \{1, \dots, G\}$,

$$\hat{u}_g = \frac{1}{n} \sum_{i=1}^{n} (\mathbf{y}_i - \hat{\mathbf{y}}_i^{(g)})^T \mathbf{W}_i (\mathbf{y}_i - \hat{\mathbf{y}}_i^{(g)}) \tag{9}$$

to quantify the importance of the $g$th x-variable. Here,

$$\boldsymbol{W}_i = \frac{1}{n_i} \hat{\boldsymbol{V}}_i^{-\frac{1}{2}} \boldsymbol{R}_i^{-1}(\hat{\boldsymbol{\phi}}) \hat{\boldsymbol{V}}_i^{-\frac{1}{2}} \tag{10}$$

where $\hat{\boldsymbol{V}}_i$ is the $n_i \times n_i$ diagonal matrix consisting of the dose-varying variance

$$\hat{\boldsymbol{V}}_i = \begin{bmatrix} \hat{V}(d_{i1}) & 0 & \dots & 0 \\ 0 & \hat{V}(d_{i2}) & \dots & 0 \\ \vdots & \vdots & \ddots & \vdots \\ 0 & 0 & \dots & \hat{V}(d_{in_i}) \end{bmatrix} \tag{11}$$

and $\boldsymbol{R}_i(\boldsymbol{\phi}) = (R_{jk})$ the $n_i \times n_i$ working correlation matrix for the $i^{th}$ subject. By $\boldsymbol{\phi}$, we denoted the $s \times 1$ vector that fully characterises the correlation structure. The estimate of $\boldsymbol{\phi}$, $\hat{\boldsymbol{\phi}}$, was obtained by taking the moment estimators for the parameters $\boldsymbol{\phi}$ in the correlation structure based on the residuals obtained from fitting the following model

$$y_{ij} = \sum_{k=0}^{p} z_{ik} \beta_k(d_{ij}) + \varepsilon_{ij} \text{ where } i = 1, \dots, n, j = 1, \dots, n_i. \tag{12}$$

The variance function $V(d)$ in Eq (11) was estimated using techniques similar to [32].

After having obtained $\{\hat{u}_g : \ g = 1, \ldots, G\}$, we sorted gene utilities in an increasing order. That is because smaller $\hat{u}_g$ values indicate stronger marginal associations. The $x$-predictors included in the screened submodel are, then, given by

$$\widehat{\mathscr{M}}_{\tau_n} = \{1 \leq g \leq G : \ \hat{u}_g \text{ ranks among the first } \tau_n(\nu)\} \tag{13}$$

where $\tau_n(\nu)$ corresponds to the size of the submodel which is chosen to be smaller than the sample size $n$.

#### Variable selection using a group SCAD (gSCAD) penalty

Screening algorithms aim to discard all unimportant variables but tend to be conservative. In order to preserve only the most important $x$-predictors in the final model, we considered a model including the first $\tau_n(\nu)$ outranked genes and we applied a gSCAD penalty by minimising the following criterion:

$$\frac{1}{2}\sum_{i=1}^{n}\sum_{j=1}^{n_i} w_{ij}\Bigg\{ y_{ij} - \sum_{k=0}^{p} z_{ik}\sum_{l=1}^{L_k}\zeta_{lk}B_{lk}(d_{ij}) - \tag{14}$$

$$\sum_{g\in\widehat{\mathscr{M}}_{\tau_n}} x_{ig}\sum_{l=1}^{L_g}\eta_{lg}B'_{lg}(d_{ij})\Bigg\}^2 + \sum_{g\in\widehat{\mathscr{M}}_{\tau_n}} p_{\lambda,\alpha}(||\boldsymbol{\eta}_g||) \tag{15}$$

where

$$p_{\lambda,\alpha}(u) = \begin{cases} \lambda u & \text{if } 0 \leq u \leq \lambda \\ -\frac{(u^2 - 2\alpha\lambda u + \lambda^2)}{2(\alpha-1)} & \text{if } \lambda \leq u \leq \alpha\lambda \\ \frac{(\alpha+1)\lambda^2}{2} & \text{if } u \geq \alpha\lambda, \end{cases}$$

$\alpha$ is a scale parameter, $\lambda$ controls for the penalty size and $||\cdot||$ is the Euclidean $L_2$-norm. At this point, note that grouping is applied for the coefficients $\boldsymbol{\eta}_g$ that correspond to the same coefficient function. In addition, in order to reduce the bias introduced when applied a LASSO penalty, we alternatively chose the SCAD, which coincides with the LASSO until $u = \lambda$, then transits to a quadratic function until

$u = \alpha\lambda$ and then it remains constant $\forall u > \alpha\lambda$, meaning that retains the penalisation and bias rates of the LASSO for small coefficients but at the same time relaxes the rate of penalisation as the absolute value of the coefficients increases. In Fig 1 the reader can find a brief overview of the employed methodology.

**Fig 1. The two-stage algorithm for identifying dose-dependent associations between genes and drugs. Gene expression and drug response data from a drug screening study (e.g. GDSC) are used to fit our dose-varying coefficients model to estimate the dose-varying effect between covariates and drug response. A two-stage variable screening and selection algorithm is applied to rank gene-drug associations. The selected genes can then be used to predict dose-dependent response for drugs of interest.**

### Tuning parameters selection

We used knots placed at the median of the observed data values along with cubic B-splines with 1 interior knot, resulting from calculating the number of interior knots suitable using the formula $N_n = [n^{\frac{1}{2p+3}}]$ proposed and applied by [29,35,36]. Due to the computational burden this would add, we did not apply cross-validation.

As for the screening threshold $\tau_n$, its magnitude could be determined by the fraction $\nu[\frac{n}{log(n)}]$, $\nu \in \{1, 2, 3, \ldots\}$. We conducted a pilot simulation study in order to decide the most appropriate size (for further details see S1 Text). We also considered an automated algorithm for its selection (Greedy Iterative Non-parametric Independence Screening-Greedy INIS, [37]). Finally, the penalty size for the gSCAD step $\lambda$ was determined using a 5-fold cross-validation.

### Simulation study

Monte Carlo simulations were conducted to examine the ability of our model to detect the genes that are truly associated with the drug response. Responses over different dosage levels were generated based on a subset of genes, the corresponding low-dimensional GDSC data covariates (drug and cancer type) and some specified smooth coefficient functions (see S1 Text). Due to the computational burden associated with a simulation of the same scale as the data set, we conducted a simulation study using smaller random fragments of the original GDSC data set. In particular, we repeatedly sampled without replacement 190 experimental units and 886 genes based on which the simulated responses have been generated. The performance of the employed methodology has been assessed based on 1000 simulations using three screening thresholds ($\tau_n(\nu) = [\frac{n}{log(n)}]$, $\tau_n(\nu) = [\frac{2n}{log(n)}]$ and $\tau_n(\nu)$ chosen using the greedy-INIS algorithm [37]) and two estimated covariance structure scenarios (independence and rational quadratic covariance structure). Cubic B-splines and knots placed at the median of the observed data values have been used for estimating the coefficient functions.

To evaluate the performance of the proposed procedure we used the following summary measures: TP−number of genes correctly identified as active; FP−number of the genes incorrectly identified as active; TN−number of the genes correctly identified as inactive; FN−number of the genes incorrectly identified as inactive.

Simulation results suggested that our method accurately detects the drug associated genes from the simulated responses under most of the examined scenarios (Fig. A in S1 Text). A screening threshold of size $[\frac{2n}{log(n)}]$ and regression weights adjusted for the covariance structure of the data have been identified as the scenario where our method reached its maximum accuracy. Consequently, for the GDSC application, we chose the screening threshold to be the maximum possible, i.e. 923 genes derived from the formula $[\frac{2n}{log(n)}]$, and weights derived by assuming a rational quadratic covariance structure for the repeated measures.

Since the *BRAF* gene is the target of the drugs, mean fold change and

**Table 1. Top 30 gene rankings based on the estimated area under the coefficient function curve.**

| Gene Name | Area | SD | Sign | Spearman's Correlation | Mean fold change in *BRAF* mutant vs wild-type cell lines | Protein-protein interaction network distance to *BRAF* |
|---|---|---|---|---|---|---|
| *KIR3DL1* | 0.370 | 0.107 | - | -0.874 | 0.978 | 3 |
| *CHST11* | 0.257 | 0.092 | - | -0.817 | 0.899 | NI |
| *APOC1P1* | 0.247 | 0.09 | - | -0.918 | 1.190 | NI |
| *PLEKHA6* | 0.239 | 0.086 | - | -0.908 | 1.037 | 3 |
| *PPM1F* | 0.223 | 0.068 | + | 0.910 | 0.883 | 3 |
| *BFSP1* | 0.222 | 0.074 | - | -0.800 | 1.217 | NI |
| *PPP1R3A* | 0.217 | 0.082 | + | 0.774 | 1.078 | 3 |
| *C16orf87* | 0.207 | 0.087 | + | 0.851 | 0.977 | NI |
| *PARVA* | 0.203 | 0.081 | + | 0.890 | 0.984 | 2 |
| *SLC39A13* | 0.202 | 0.079 | - | -0.461 | 1.055 | NI |
| *UCN2* | 0.198 | 0.07 | - | -0.928 | 0.979 | NI |
| *STMN3* | 0.198 | 0.087 | + | 0.834 | 1.201 | 2 |
| *RNF130* | 0.197 | 0.083 | - | -0.927 | 1.153 | NI |
| *C3orf58* | 0.196 | 0.076 | + | 0.922 | 1.133 | NI |
| *CXXC4* | 0.188 | 0.079 | + | 0.866 | 0.995 | NI |
| *THBD* | 0.179 | 0.093 | 0 | -0.967 | 1.231 | 4 |
| *SIRT3* | 0.173 | 0.066 | - | -0.760 | 1.013 | 3 |
| *PLAT* | 0.172 | 0.092 | - | -0.878 | 1.322 | 4 |
| *MPPED1* | 0.168 | 0.066 | + | 0.430 | 0.978 | NI |
| *INSL3* | 0.162 | 0.068 | - | -0.973 | 0.965 | NI |
| *FAM163A* | 0.159 | 0.078 | - | -0.983 | 1.106 | NI |
| *CNIH3* | 0.153 | 0.08 | - | -0.918 | 0.938 | NI |
| *GJA3* | 0.153 | 0.067 | 0 | -0.940 | 0.933 | NI |
| *BTG2* | 0.152 | 0.078 | + | 0.959 | 1.035 | 2 |
| *DLX6* | 0.152 | 0.059 | 0 | 0.686 | 0.987 | NI |
| *DLC1* | 0.151 | 0.053 | - | -0.928 | 0.974 | 3 |
| *GAPDHS* | 0.150 | 0.077 | + | 0.886 | 1.232 | NI |
| *JAG2* | 0.149 | 0.069 | - | -0.994 | 0.981 | 3 |
| *SMOX* | 0.146 | 0.057 | 0 | 0.816 | 1.070 | NI |
| *ZMYND8* | 0.145 | 0.091 | + | 0.907 | 1.020 | 3 |

**Table 2. Rankings of the genes identified from the pathway and oncogenic gene set enrichment analysis.**

| Gene Name | Area | SD | Sign | Spearman's Correlation | Mean fold change in *BRAF* mutant vs wild-type cell lines | Protein-protein interaction network distance to BRAF |
|---|---|---|---|---|---|---|
| MYO5A | 0.531 | 0.261 | + | 0.955 | 1.358 | 4 |
| S100A1 | 0.488 | 0.189 | + | 0.812 | 1.263 | NI |
| GPNMB | 0.424 | 0.196 | + | 1 | 1.169 | 3 |
| ACP5 | 0.359 | 0.149 | - | -0.998 | 1.039 | NI |
| FCGR2A | 0.341 | 0.158 | - | -0.588 | 1.25 | 3 |
| CITED1 | 0.28 | 0.348 | 0 | -0.603 | 1.63 | 3 |
| SPRY4 | 0.274 | 0.127 | - | -0.611 | 1.228 | 2 |
| CD44 | 0.239 | 0.164 | + | 0.868 | 1.413 | 3 |
| RAP2B | 0.236 | 0.179 | 0 | 0.927 | 1.254 | NI |
| KCNJ13 | 0.205 | 0.094 | 0 | -0.604 | 1.101 | 3 |
| ALX1 | 0.202 | 0.099 | - | -1 | 1.104 | NI |
| PLAT | 0.201 | 0.121 | - | -0.405 | 1.312 | 4 |
| RETSAT | 0.201 | 0.142 | 0 | 0.689 | 1.127 | NI |
| GSN | 0.196 | 0.109 | + | 0.588 | 1.079 | 4 |
| CDH19 | 0.185 | 0.102 | 0 | 0.943 | 0.933 | NI |
| ATP1B3 | 0.178 | 0.115 | - | -1 | 1.063 | NI |
| BAZ1A | 0.173 | 0.105 | + | -0.29 | 1.109 | 4 |
| SLC16A4 | 0.166 | 0.117 | - | -0.298 | 1.234 | NI |
| ST6GALNAC2 | 0.164 | 0.102 | 0 | -0.815 | 1.264 | NI |
| MFSD12 | 0.16 | 0.148 | 0 | -0.788 | 1.13 | NI |
| GJA3 | 0.157 | 0.075 | 0 | -0.85 | 1.071 | NI |
| CYP27A1 | 0.156 | 0.09 | - | -0.743 | 1.373 | NI |
| EGLN1 | 0.15 | 0.119 | - | -0.442 | 1.053 | 3 |
| TRPV2 | 0.147 | 0.118 | 0 | 0.769 | 1.074 | NI |
| MITF | 0.146 | 0.106 | + | 1 | 0.743 | 2 |
| TBC1D7 | 0.146 | 0.118 | 0 | -0.603 | 1.304 | NI |
| SLC6A8 | 0.144 | 0.111 | 0 | -0.263 | 0.941 | NI |
| PTPRZ1 | 0.139 | 0.138 | - | -0.808 | 1.074 | 4 |
| PLOD3 | 0.132 | 0.135 | 0 | 0.696 | 1.166 | NI |
| ANKRD7 | 0.131 | 0.12 | + | 0.92 | 1.241 | NI |
| KANK1 | 0.107 | 0.113 | 0 | -0.493 | 1.345 | NI |
| GYPC | 0.105 | 0.092 | + | -0.3 | 1.072 | NI |
| TYR | 0.1 | 0.098 | - | 0.467 | 1.11 | 4 |
| TYRP1 | 0.1 | 0.097 | 0 | 0.457 | 1.326 | 3 |
| IGSF8 | 0.09 | 0.129 | 0 | -0.668 | 1.313 | 5 |
| SPRED1 | 0.067 | 0.116 | 0 | -0.556 | 1.239 | 4 |
| ITGA9 | 0.056 | 0.111 | 0 | 0.785 | 1.154 | 4 |
| KREMEN1 | 0.053 | 0.086 | 0 | -0.555 | 1.123 | 4 |
| LAMA4 | 0.038 | 0.083 | - | 0.344 | 1.151 | 4 |
| MLANA | 0.037 | 0.097 | 0 | 0.534 | 1.147 | NI |
| KLF9 | 0.011 | 0.074 | 0 | 0.932 | 1.064 | NI |

Table notes rankings of the genes found to have some biological importance. A positive (+) sign translates to a positive effect on cells survival after drug administration, a negative (-) sign translates to a negative effect on cells survival and a neutral (0) effect translates to a varying effect on cells survival which depends on drug dosage. Spearman's correlation is calculated between drug dosage and gene estimated coefficient function values as an indicator of the magnitude change of the gene effect over the increasing dosage. Area corresponds to the area under the estimated coefficient curve and the SD corresponds to the standard deviation of the area based on bootstrapping. Pearson's correlation is calculated between the selected gene microarray expression values and the *BRAF* expression across all the cell lines. Protein-protein interaction network distance is computed based on the shortest interaction path between the *BRAF* gene and each of the selected genes. Here, NI denotes absence of interaction.

comes to predicting the most effective drug or dosage range (≈79% in both scenarios). Results are less reliable when it comes to prediction of the exact dosage or drug-dosage combination (≈48-49% and ≈57-58% in both scenarios) but this can be due to either the large variability observed in the observed responses or due to the small number of cell lines for some predictor level combinations. Results were similar for both cross-validation scenarios (differences range from 0 to <2%, Table 3), meaning that as long as a cell line has similar genetic characteristics to those observed, the model can be reliable in predicting the outcome after anticancer drug administration.

**Table 3. Predictive performance of the employed model (mean absolute error=0.121).**

| **Scenario** | **Accuracy EU** | Accuracy CL |
|---|---|---|
| Model predicts the more effective drug-dosage combination | 57.85% | 57.42% |
| Model predicts the more effective drug given a cell-line | 78.21% | 78.21% |
| Model predicts the more effective dosage given a drug | 48.44% | 48.65% |
| Model predicts the most effective dosage range ($>$ or $\leq$ 31.25% of the maximum dosage) | 79.47% | 79.28% |

Table notes the predictive performance of the model based on the percentages of correctly identifying the most effective drug, dosage or drug-dosage combinations. Results obtained based on 10-fold cross-validation of the final model (based on holding out either experimental units−EU− or cancer cell lines−CL−).

Drug response prediction from gene expression data has been widely studied in the literature. Sparse regression methods, gene selection algorithms such as the Ping-pong algorithm [25], or a combination of network analysis and penalised regression, e.g. the sparse network-regularized partial least squares method [17], have all been employed to simultaneously predict drug response and select genetic factors that seem to be associated with the drug response. However, none of these methods are able to quantify the effect of drug dosage on the response. Employing the proposed dose-varying model gives a detailed picture of different drugs effect and can be extremely valuable in predicting drug response for agents with small therapeutic range and high toxicity levels. In addition, the method can be easily extended for different cell lines-drug combinations as well as different types of molecular data (e.g. RNA-seq gene expression, methylation or mutational profiles). Finally, due to the structure of our model, enrichment with additional low-dimensional covariates, such as drug chemical information, is straightforward.

50. Subramanian A, Tamayo P, Mootha VK, Mukherjee S, Ebert BL, Gillette MA, et al. Gene set enrichment analysis: a knowledge-based approach for interpreting

genome-wide expression profiles. Proceedings of the National Academy of Sciences. 2005;102(43):15545–15550.

51. Benjamini Y, Hochberg Y. Controlling the false discovery rate: a practical and powerful approach to multiple testing. Journal of the Royal statistical society: series B (Methodological). 1995;57(1):289–300.

52. Solit DB, Rosen N. Resistance to BRAF inhibition in melanomas. New England Journal of Medicine. 2011;364(8):772–774.

53. Manzano JL, Layos L, Bugés C, de los Llanos Gil M, Vila L, Martinez-Balibrea E, et al. Resistant mechanisms to BRAF inhibitors in melanoma. Annals of translational medicine. 2016;4(12).

### Supporting information captions
